# Ureter single-cell and spatial mapping reveal cell types, architecture, and signaling networks

**DOI:** 10.1101/2021.12.22.473889

**Authors:** Emily E. Fink, Surbhi Sona, Uyen Tran, Pierre-Emmanuel Desprez, Matthew Bradley, Hong Qiu, Mohamed Eltemamy, Alvin Wee, Madison Wolkov, Marlo Nicolas, Booki Min, Georges-Pascal Haber, Oliver Wessely, Byron H. Lee, Angela H. Ting

## Abstract

Tissue engineering offers a promising treatment strategy for ureteral strictures, but its success requires an in-depth understanding of the architecture, cellular heterogeneity, and signaling pathways underlying tissue regeneration. Here we define and spatially map cell populations within the human ureter using single-cell RNA sequencing, spatial gene expression, and immunofluorescence approaches. We focused on the stromal and urothelial cell populations to enumerate distinct cell types composing the human ureter and inferred potential cell-cell communication networks underpinning the bi-directional crosstalk between these compartments. Furthermore, we analyzed and experimentally validated the importance of Sonic Hedgehog (SHH) signaling pathway in adult stem cell maintenance. The *SHH*-expressing basal cells supported organoid generation *in vitro* and accurately predicted the differentiation trajectory from basal stem cells to terminally differentiated umbrella cells. Our results highlight essential processes involved in adult ureter tissue homeostasis and provide a blueprint for guiding ureter tissue engineering.

## Introduction

The ureter is a tube that carries urine from the kidney to the bladder; it is often overshadowed by the two organs that it connects and neglected in molecular studies. However, congenital defects and injuries of this structure can lead to significant medical issues, which severely compromise patients’ quality of life. Ureteral stricture is a common sequelae of iatrogenic injury from surgery or radiation but can also be caused by benign conditions such as impacted ureteral calculus, retroperitoneal fibrosis, or abdominal aortic aneurysm (Gild et al., 2018; Vorobev et al., 2021). If left untreated, ureteral stricture can lead to pain, repeat infections, kidney stones, and permanent loss of renal function. The surgical repair approach depends on stricture location, length, density, and other factors such as previous treatments and/or radiation. Long ureteral strictures can be particularly challenging to repair and can involve the use of non-native tissue such as buccal mucosal graft, appendiceal flap, or ileal ureteral interposition (Gild et al., 2018; Paffenholz and Heidenreich, 2021). Long-term complications of bowel use in the urinary tract include recurrent urinary tract infection (UTI), metabolic derangements, and potential renal function deterioration (Gild et al., 2018).

Tissue engineering of human ureters is an emerging field and could be an alternative treatment approach in these situations. However, for successful ureter tissue engineering, several problems need to be considered, including physiological characteristics and local environment of the tissue, the type of scaffold used, and, most importantly, a detailed understanding of the individual cell types in the adult human ureter (Simaioforidis et al., 2013). Advancements in single-cell RNA sequencing technologies have revolutionized our understanding of the cellular complexity for a myriad of different tissue types, both in normal and diseased states (Wolfien et al., 2021). Single cell transcriptomics has been used to study the mouse and human kidney (Combes et al., 2019; Lake et al., 2019; Menon et al., 2018; Sheng et al., 2021; Wu et al., 2018); however, these analyses have focused on interrogating the cellular composition of the kidney with a focus on the kidney interstitium and nephrons, but have not described the urothelial compartment. In fact, with respect to the urothelium, single cell transcriptomic studies have been reported only on the human bladder (Chen et al., 2020; Li et al., 2012; Yang et al., 2017; Yu et al., 2019), but not the human ureter. Historically, the urothelium of the ureter has been overlooked and/or considered continuous with the urothelium of the bladder. While they share many characteristics, the ureter and bladder are fundamentally different tissues with the ureter arising from the intermediate mesoderm and the bladder arising from the endoderm (Rehman and Ahmed, 2021). Therefore, an in-depth analysis of the human ureter is critical to uncover and understand novel developmental networks and physiological aspects of this tissue.

In this study, we defined the single-cell landscape of human ureter tissues comprised of 30,942 cells from ten different patients. Analysis of these data revealed a previously underappreciated complexity in the immune, stromal and urothelial compartment. We identified a proliferative stem-like basal urothelial cell population, with potential functional roles in response to tissue injury/inflammation, as well as the ability to support the *in vitro* generation of organoids. We also investigated the detailed spatial relationships among four types of fibroblasts by complementing single-cell transcriptomic data with spatial transcriptomics and immunofluorescence.

Furthermore, we interrogated potential cellular communications between fibroblasts and urothelial cells, enhancing our understanding of the putative pathways involved in homeostasis and regeneration. These results promote a detailed understanding of the cellular heterogeneity and signaling networks within the human ureter and serve as the foundation for future tissue engineering endeavors.

## Results

### Single-cell profiling and unbiased clustering of human ureter cells

To uncover the cellular complexity of the human ureter, we performed single-cell RNA sequencing (scRNAseq) on human ureter tissues. Normal human ureter tissue was procured from patients undergoing radical cystectomy, immediately dissociated into single cell suspensions, and processed for scRNAseq. After quality control and filtering, a total of 30,942 single cells from ten different human ureter samples were segregated into 20 different clusters using Seurat’s unsupervised clustering algorithm (**Tables S1** and **S2**). These clusters were visualized by Uniform Manifold Approximation and Projection (UMAP), where each color represents a different cell population ordered from the largest (4,200 cells in cluster 0) to the smallest (94 cells in cluster 19) (**Figure 1A**). Patient samples and sexes were uniformly distributed throughout the UMAP with the exception of cluster 5, which was dominated by female samples, with the highest proportion of cells coming from sample U7 (**Figures S1A** and **S1B**). We then performed differential gene expression analysis to aid in the classification of each cell cluster (**Figures 1B** and **S1C**; **Table S2**). Leveraging established markers, we could separate the clusters into three major compartments, urothelial (eight clusters expressing *KRT20*, *KRT8*, *KRT17*, *MIK67*, *KRT5*, *KRT13*, *UPK3A*), stromal (four clusters expressing *ACTA2*, *FLT1*, *COL1A2*, *DCN*), and immune (eight clusters expressing *TRAC*, *CD8A*, *NKG7*, *TPSB2*, *CD163*, *FCN1*, *LST1*). We noted an absence of neuronal cell types in our dataset, but this is consistent with other reports describing difficulties in isolating intact neurons during tissue dissociation (Armand et al., 2021). Pearson correlation coefficient based on the expression of the top 2,000 variable genes between all possible cluster pairs confirmed that clusters within each compartment share more similar expression profiles when compared to clusters in other compartments (**Figure 1C**), corroborating our classification. We thus proceeded to examine each subset separately to improve the resolution of cell types.

**Figure 1.**
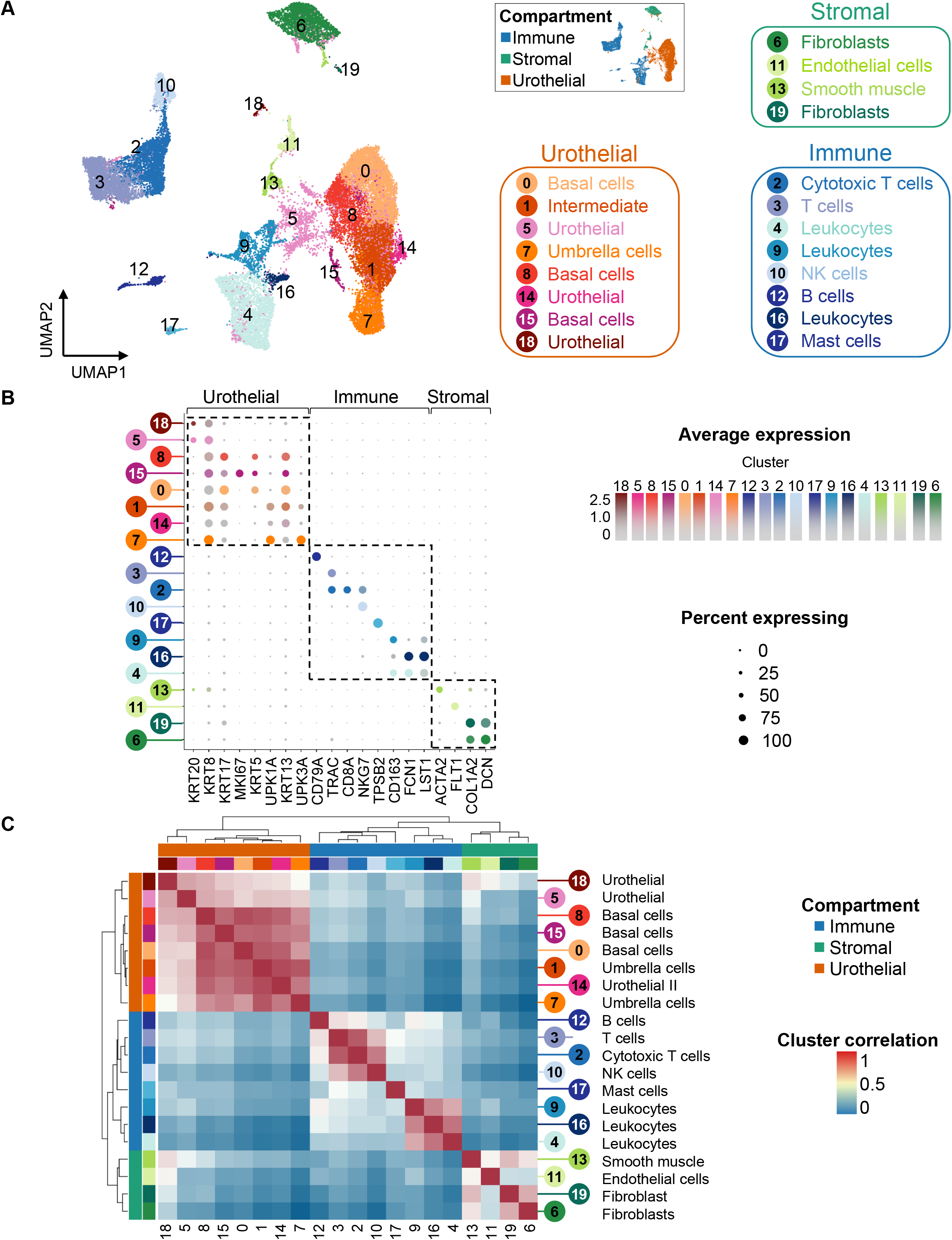
Single-cell profiling and unbiased clustering of human ureter cells. (A) UMAP based on gene expression. Cells are colored by cluster. Clusters belong to 3 major tissue compartments (inset) – immune (blue), stromal (green) and urothelial (orange). (B) Dot plot shows the expression level of known markers across cell cluster. (C) Pearson’s correlation matrix of the average expression profiles of each cluster, based on the top 2000 variable genes. See also Figure S1 and Tables S1-S2.

### Classification of the immune cells

Reanalysis of the immune subset resulted in 16 clusters, with minimal patient-specific and sex-specific effects (**Figures S2A** and **S2B**; **Table S3**). Following differential gene expression analysis, we identified one cluster each for mast cells (cluster 13: *KIT*, *TPSAB1*), NK cells (cluster 3: *KLRD1*, *CD247*), and B cells (cluster 6: *CD79A*, *JCHAIN*) (**Figure 2A**). We also observed several clusters within the monocyte lineage (clusters 0, 7, 8, 10, and 12) grouped together on the UMAP, with varying expression of the monocyte marker *CD14* (Zamani et al., 2013). Higher levels of *CD14* were observed in dendritic cells (cluster 7: *HLA-DRA*, *HLA-DRB1*) and macrophages (cluster 8: *CD14*, *C1QA*) (**Figures 2B** and **S2C**). Three *CD14*/*CD68*-expressing monocyte clusters with high expression of ficolin 1 (*FCN1*) were also identified, and they were further divided into classical (clusters 0 & 10: *S100A8, S100A9*) and non-classical (cluster 12: *MS4A7, LILRB2, SELPLG*) monocytes (Hu et al., 2020). Interestingly, cluster 0 classical monocytes appeared to co-express intermediate markers, including *BCL6* and *STAT3* (Loperena et al., 2018).

**Figure 2.**
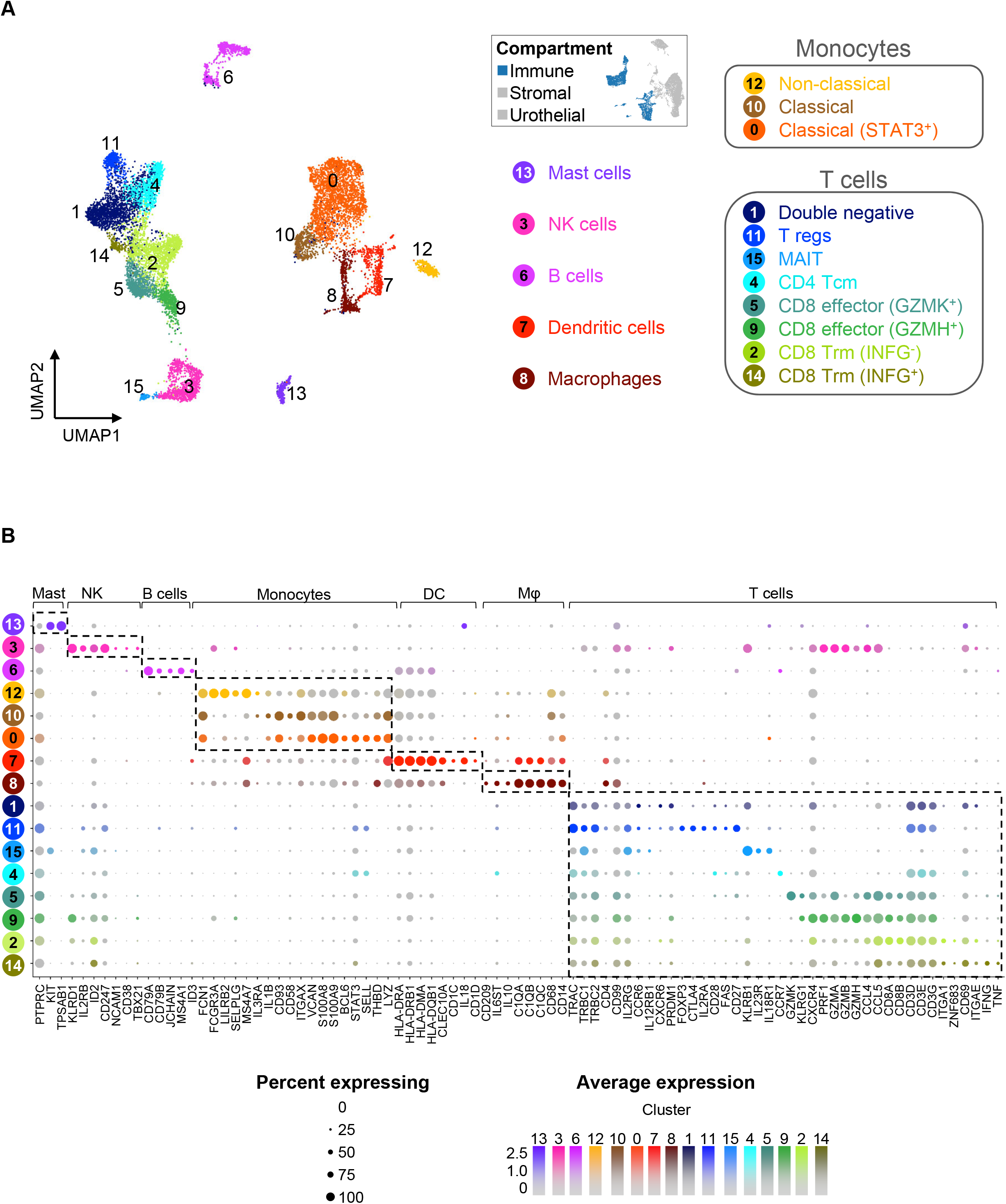
Classification of the immune cells based on cell type-specific marker genes. (A) UMAP based on gene expression. Cells are colored by cluster. (B) Dot plot shows the expression level of known immune cell markers across cell clusters. See also Figure S2 and Table S3.

We also identified eight different clusters of T cells (clusters 1, 2, 4, 5, 9, 11, 14 & 15). The largest T cell cluster was negative for *CD4* and *CD8* expression and was referred to as double negative T cells (cluster 1). Also amongst the T cell clusters were regulatory T (Treg) cells (cluster 11: *FOXP3*, *CTLA4*), central memory T (Tcm) cells (cluster 4: *CCR7*), and two clusters of tissue resident memory T (Trm) cells (clusters 2 & 14: *ZNF683*, *CD69*, *ITGA1*). Expression of cytokines *INFG* and *TNF* in cluster 14 Trm cells suggests that this cluster may have been recently activated. We also identified two clusters of CD8 effector T cells, the *GZMK*+ cluster 5 and the *GZMH*+ cluster 9. Finally, we resolved a cluster of mucosal-associated invariant T (MAIT) cells (cluster 15: *KLRB1*, *IL23R*, *IL18R1*), which are innate-like T cells that play an important role in the antibacterial response (Godfrey et al., 2019; Terpstra et al., 2020).

It is important to note that the ten ureter tissues used for the single cell study were free from any existing injury or infection, based on medical record and pathologist examination of the tissues, with the potential exception of patient U7, who had a history of chronic urinary tract infections (UTIs). Furthermore, in the urinary tract, it has been reported that the adaptive immune response is limited, requiring an efficiently primed innate immune component (Hayes and Abraham, 2016). Therefore, the monocyte cell types (clusters 0, 10, and 12) likely play important homeostatic roles (Teh et al., 2019). Nonetheless, the heterogeneity of immune cell types in normal human ureters was strikingly high.

### Classification of the stromal cells

The analysis of the stromal cell subset resulted in seven clusters (**Figure 3A**), with even distribution across patient samples and sexes (**Figures S3A** and **S3B**). Each stromal cluster expressed a unique gene set (**Figure S3C**; **Table S4**), allowing for unambiguous annotation of each cell type. As expected, we identified two *PECAM1*^+^ endothelial cell clusters, which could be separated into venous endothelial cells (cluster 1: *NRP1*, *NRP2*), and arterial endothelial cells (cluster 6: *GJA5*, *BMX*) (**Figure 3B**) (dela Paz and D’Amore, 2009). We also identified one smooth muscle cell cluster (cluster 2: *ACTA2*, *MYH11*) and four distinct fibroblast clusters (clusters 0, 3, 4, and 5). Cluster 5 fibroblasts expressed high levels of *COL1A1* and was the most distinct from the other fibroblasts in the ureter based on its position on the UMAP. The others, while clustered closely together, could still be clearly distinguished by unique markers (**Figure S3C**) and were labeled *HAS1*^hi^ (cluster 0), *APOE*^hi^ (cluster 3), and *GAS1*^hi^ (cluster 4).

**Figure 3.**
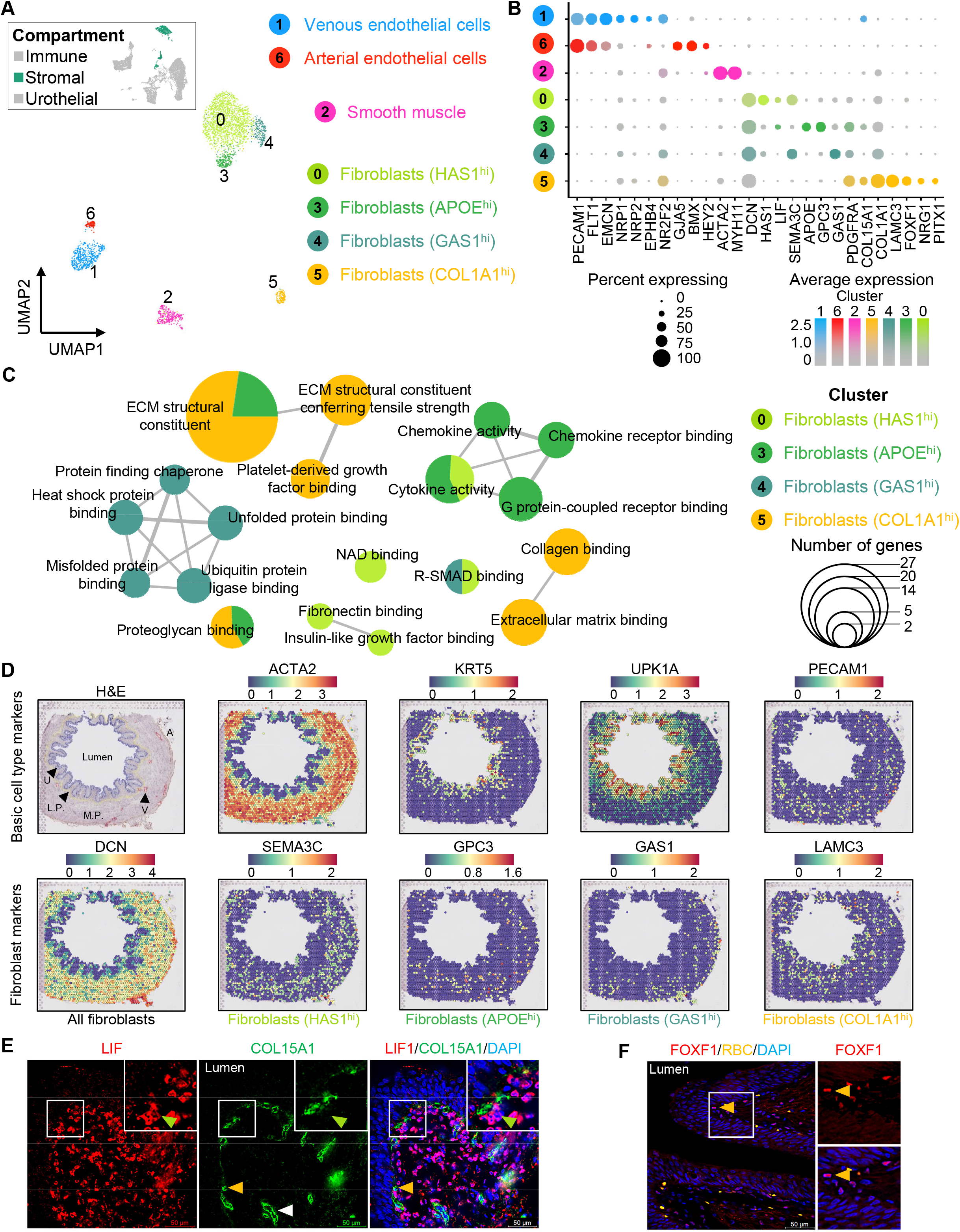
Classification of the stromal cells based on cell type-specific marker genes. (A) UMAP based on gene expression. Cells are colored by cluster. (B) Dot plot shows the expression level of known markers across cell clusters. (C) Comparison of the 4 fibroblast clusters using enrichment analysis performed on upregulated differential fibroblast markers. (D) Visium spatial gene expression in log-normalized (logNorm) counts for selected genes in human ureter cross-sections. The H&E image is annotated with features for orientation: lumen, urothelium (U), lamina propria (L.P.), muscularis propria (M.P.), blood vessel (V), and adventitia (A). (E) & (F) Immunofluorescence staining of human ureter cross-sections. Arrowheads are color-coded by cluster. The white arrowhead in the COL15A1 panel identifies blood vessels. See also Figure S3 and Table S4

To gain insight into the functional differences between the four fibroblast clusters, we performed gene set enrichment analysis on differentially expressed genes across the fibroblasts (**Figure 3C**). Each fibroblast cluster had unique Gene Ontology (GO) term enrichments, suggesting that they were functionally distinct. Cluster 5 fibroblasts (*COL1A1^hi^*) likely serve a prominent structural role and were significantly enriched for GO terms related to extracellular matrix binding and collagen binding. Cluster 3 fibroblasts (*APOE^hi^*) were enriched for terms related to chemokine activity, and in the skin, apolipoprotein E (*APOE*) expressing fibroblasts were thought to have pro-inflammatory properties (Sole-Boldo et al., 2020). Cluster 4 fibroblasts (*GAS1^hi^*) were enriched for terms related to unfolded protein and heat shock protein binding. In fact, growth arrest specific-1 (*GAS1*) is an anti-mitogenic factor that mediates growth arrest of fibroblasts. The *Saccharomyces cerevisiae* homologue of *GAS1* plays an important role in the endoplasmic reticulum stress response (Cui et al., 2019). Thus, this population of fibroblasts might play a role in upregulating pathways involved in protein homeostasis. The largest cluster of fibroblasts, cluster 0 (*HAS1*^hi^), was enriched for the term “fibronectin binding”. Hyaluronan synthase 1 (*HAS1*) is responsible for the production of hyaluronan, which is a key constituent of the extracellular matrix and is involved in cell migration, wound healing, and tissue repair (Jiang et al., 2007). Hyaluronan has also been implicated in regulating the deposition of fibronectin and collagen (Evanko et al., 2015), which together provide the framework for the ingrowth of blood vessels and fibroblasts during tissue healing and regeneration.

In a different attempt to better understand the four fibroblast populations, we leveraged the Visium Spatial Gene Expression platform to generate spatial transcriptomic profiles of human ureter cross sections. While this analysis did not reach single-cell resolution, it allowed us to clearly distinguish the urothelial layer (*KRT5*^+^ basal cells and *UPK1A*^+^ umbrella cells) from the outer stromal layer (*ACTA2*^+^ smooth muscle cells and *DCN*^+^ fibroblasts). Therefore, by querying the markers used to specify each fibroblast cluster in the scRNAseq analysis, we could visualize how each fibroblast subtype was distributed in the stromal layer. The markers chosen were *SEMA3C*, *GPC3*, *GAS1*, and *LAMC3* for clusters 0, 3, 4, and 5, respectively (**Figures 3D** and **S3D**). Clusters 0 (*HAS1*^hi^) and 3 (*APOE*^hi^) were diffusely spread throughout the interstitial layer. Cluster 4 (*GAS1*^hi^) was localized towards the periphery of the interstitial layer. Finally, Cluster 5 (*COL1A1*^hi^) was localized closest to the urothelium in contrast to the other fibroblast clusters.

Finally, we used commercially available antibodies that specifically distinguished, either alone or in combination, the fibroblast subtypes. Cluster 0 was identified by being LIF1^+^/COL15A1^-^ and Cluster 5 by LIF1^-^/COL15A1^+^ (**Figure 3E**). Cluster 5 could also be uniquely visualized as FOXF1^+^ cells, which are localized just below the urothelium and is consistent with the spatial gene expression data (**Figure 3F**). Moreover, cluster 3 fibroblasts clearly stained for APOE and could be seen interspersed between the ACTA2^+^ smooth muscle bundles (**Figure S3E**). The immunofluorescence staining not only validated the RNA expression-based classification of the fibroblasts but also confirmed their spatial distributions within the tissue. Together, these data demonstrate a so far unappreciated heterogeneity of the ureter interstitium that, like in the kidney, could be involved in providing critical inputs for ureter homeostasis (England et al., 2020).

### Classification of the urothelial cells

The urothelial compartment could be resolved into eight clusters (**Figure 4A**), where samples and sexes were evenly distributed across clusters (**Figures S4A** and **S4B**). We identified two umbrella cell clusters (4 & 7), three intermediate cell clusters (0, 1, & 5), and three basal cell clusters (2, 3, & 6) (**Figure S4C**; **Table S5**).

**Figure 4.**
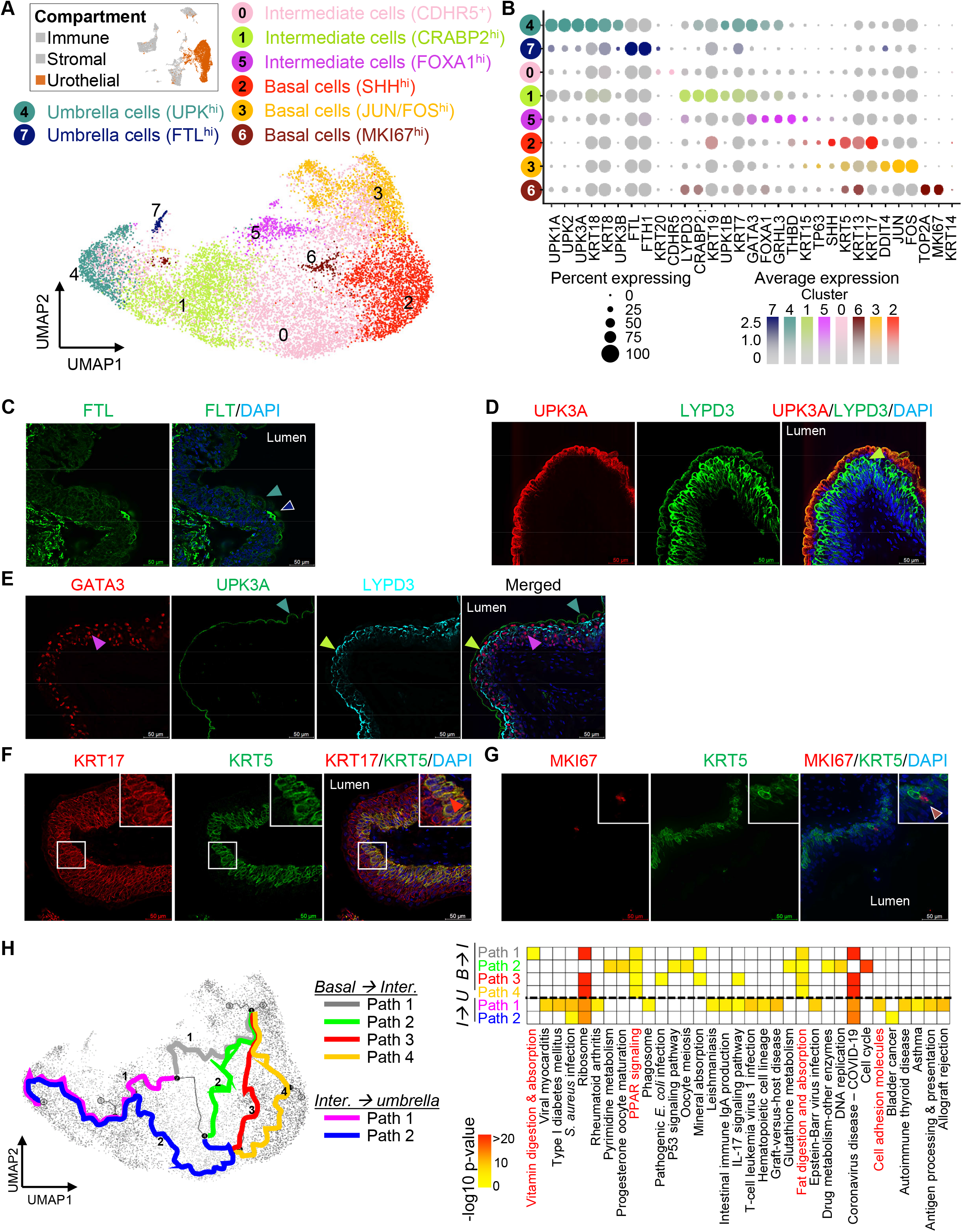
Classification of the urothelial cells based on cell type-specific marker genes. (A) UMAP based on gene expression. Cells are colored by cluster. (B) Dot plot shows the expression level of known markers across cell cluster. (C - G) Immunofluorescence staining of human ureter cross-sections. Arrows are color-coded by cluster. (H) Trajectory analysis of urothelial subset (left panel). Genes that vary the most along each path were subjected to enrichment analysis using gProfiler, and the significantly enriched pathways were compared across these different paths by their −log p value plotted on a heatmap (right panel). See also Figure S4 and Tables S5-6.

Umbrella, intermediate, and basal cells are the known major cell types within the urothelium, which span the differentiation spectrum from the least differentiated basal cells to the terminally differentiated umbrella cells (Dalghi et al., 2020). Uroplakin (*UPK*) genes are routinely used as markers for umbrella cells, but studies have shown that the umbrella cells of the ureter are more heterogeneous with respect to *UPK* expression than the bladder (Dalghi et al., 2020; Riedel et al., 2005). Cluster 4 umbrella cells were defined by high expression of *UPK* genes (*UPK1A*, *UPK2*, *UPK3A*) and labeled as *UPK*^hi^ (**Figure 4B**). Conversely, cluster 7 cells expressed low levels of *UPK* genes but showed *KRT20*, which is often used as a terminal differentiation marker (Volkmer et al., 2012). Interestingly, cluster 7 umbrella cells expressed high levels of ferritin light (*FTL*) and heavy (*FTH1*) chain genes. This was independently confirmed with immunofluorescence staining, which revealed FTL^+^ or KRT20^+^ umbrella cells as scattered in the expected abundance throughout the lumen-facing umbrella layer (**Figures 4C** and **S4D**).

We next examined the genes differentially expressed by the three intermediate cell clusters (**Figure 4B**). Cluster 0, which is the largest and most diffuse cluster, had very few distinguishing genes with the exception of cadherin-related family member 5 (*CDHR5*). Cluster 1 was defined by high expression of cellular retinoic acid-binding protein 2 (*CRABP2*) and LY6/PLAUR domain containing 3 (*LYPD3*). Cluster 5 showed high expression of *FOXA1* transcription factor, which is involved in bladder differentiation (DeGraff et al., 2012), and grainyhead like transcription factor 3 (*GRHL3*), which drives terminal differentiation and barrier formation in the urothelium (Yu et al., 2009). Interestingly, in contrast to clusters 0 and 5, cluster 1 intermediate cells showed expression of *UPK* genes, suggesting that these cells might be differentiating into the umbrella cells. Indeed, immunofluorescence analysis demonstrated that intermediate cells with the highest levels of LYPD3 were just below the UPK3A^+^ umbrella cells (**Figure 4D**). Cluster 5 intermediate cells (*FOXA1^hi^*) were located deeper in the urothelial cell layer as shown by low GATA3/LYPD3 staining pattern (**Figure 4E**).

Finally, there were three *KRT5*^+^ basal cell subtypes. Cluster 2 (*SHH^hi^*) had the highest proportion of sonic hedgehog signaling molecule (*SHH*) expressing cells (**Figure 4B**). Cluster 3 (*JUN/FOS^hi^*) also contained a small number of *SHH*-expressing cells but was saliently marked by robust expression of *JUN* and *FOS* transcription factors. Cluster 6, the smallest of the *KRT5*^+^ basal cells, was unique in expressing high levels of proliferative genes including *MKI67*, *PCLAF*, *TYMS*, *TK1*, *BIRC5*, and *RRM2* (**Figure S4C**) and was referred to as *MKI67*^hi^. The basal cell clusters could be validated by immunofluorescence staining. A gradient of KRT5 and KRT17 expression marked cluster 2 with strong double staining and cluster 3 with weaker double staining (**Figure 4F**) while cluster 6 (*MKI67*^hi^) was distinctly positive for MKI67 within the KRT5^+^ cell layer (**Figure 4G**). Previous studies have indicated KRT14^+^/KRT5^+^ basal urothelial cells as the proliferating pool of basal cells responsible for replenishing the bladder urothelium (Liu et al., 2019). However, we did not detect significant levels of *KRT14* in any of the *KRT5*^+^ basal cell clusters (**Figure 4B**). This is in agreement with studies showing that, unlike the bladder urothelium, *KRT14* is found in very low levels in the adult ureter urothelium (Bohnenpoll et al., 2017a), indicating basal cell heterogeneity between the two organs.

### Reconstructing the differentiation trajectory of the human ureter urothelial cells

It is well-accepted that cells of the basal cell layer can differentiate into intermediate and umbrella cells during development as well as during adult tissue regeneration (Dalghi et al., 2020). In the developing ureter, a SHH-FOXF1-BMP4 signaling axis has been shown to regulate cellular pathways underlying ureter elongation and differentiation (Bohnenpoll et al., 2017b; Jackson et al., 2020). In mouse bladder tissues, a SHH-expressing basal stem cell population can regenerate all the urothelial cell types following tissue injury or infection (Shin et al., 2011). Sonic hedgehog signaling molecule (SHH) is a secreted protein and a potent inducer of WNT stromal expression, stimulating proliferation of both the stromal and urothelial cells. This signaling cascade and associated proliferation are also necessary for the restoration of the bladder urothelium following injury or infection. Our scRNAseq data clearly identified a population of *SHH*-expressing basal cells in the adult ureter tissue, although the role of *SHH* in adult ureter homeostasis has not been investigated.

We began exploring the relationship among the ureter urothelial cell types with respect to SHH signaling by performing a cell trajectory analysis using Monocle3 (Cao et al., 2019; Qiu et al., 2017; Trapnell et al., 2014) and designating the cells with the highest *SHH* expression (in cluster 3) as the starting point for the trajectory (**Figure 4H**). Remarkably, this analysis recapitulated urothelial differentiation from *SHH*^hi^ basal cells, through the intermediate cells, to the terminally differentiated *UPK*-expressing umbrella cells. Interestingly, this trajectory did not exist as a single path, but instead had four paths for basal-to-intermediate cell differentiation and two paths for intermediate-to-umbrella cell differentiation. This suggested a higher degree of plasticity than previously anticipated.

To better understand how signaling pathways are changing along the differentiation paths, we performed KEGG pathway enrichment analysis of the top 200 differentially expressed genes from each differentiation trajectory (**Figure 4H**; **Table S6**). Surprisingly, only one of the basal-to-intermediate paths (#2) traversed through the *MKI67^hi^* proliferative cluster and was unique for enrichment of the KEGG pathway terms “cell cycle” and “DNA replication”. The overarching theme for the four basal-to-intermediate trajectories was enrichment in “PPAR signaling” as well as “fat digestion and absorption”. Of note, peroxisome proliferator-activated receptor (PPAR) signaling has been shown to be an important upstream regulator of urothelial differentiation in mouse bladders (Liu et al., 2019). The two intermediate-to-umbrella trajectories were not enriched for “PPAR signaling” or “fat digestion and absorption”, suggesting that these signaling networks are only critical for the early stages of differentiation. Consistent with this notion, key genes in the PPAR signaling network were highly expressed in the basal and intermediate cells but downregulated in umbrella cells (**Figures S4E** and **S4F**). Finally, the intermediate-to-umbrella trajectory 1 was enriched for “cell adhesion molecules”, signifying barrier formation. Taken together, our cell trajectory analysis showed that the manual selection of *SHH*-expressing basal cells as the starting point can accurately recapitulate the differentiation trajectory of basal cells to umbrella cells. Pathway enrichment analysis of the genes that were dynamic across the differentiation trajectories highlighted a potential role for PPAR signaling in the early stages of differentiation and a switch towards the acquisition of barrier function during terminal differentiation.

### Determining the cell-cell communication networks in the adult human ureter

The bidirectional crosstalk between the urothelium and the stroma is essential for the proper development of the urothelium and smooth muscle in both bladder and ureter (Jackson et al., 2020). To examine such communication networks between the urothelial and non-urothelial cells, we applied CellPhoneDB (Efremova et al., 2020) to infer potential receptor-ligand interactions between all cell clusters (**Table S7**). The analysis revealed *COL1A1^hi^* fibroblasts to have the most putative interactions with other stromal and urothelial cell types whereas interactions to, from, and within the immune compartment were rather muted (**Figures 5A** and **5B**). These highly interactive *COL1A1^hi^* fibroblasts reside just below the basal cell layer (**Figures 3D–F**), prompting us to examine more closely the potential signaling from this fibroblast cluster to the three basal urothelial clusters (**Figure 5C**) and vice versa (**Figure 5D**).

**Figure 5.**
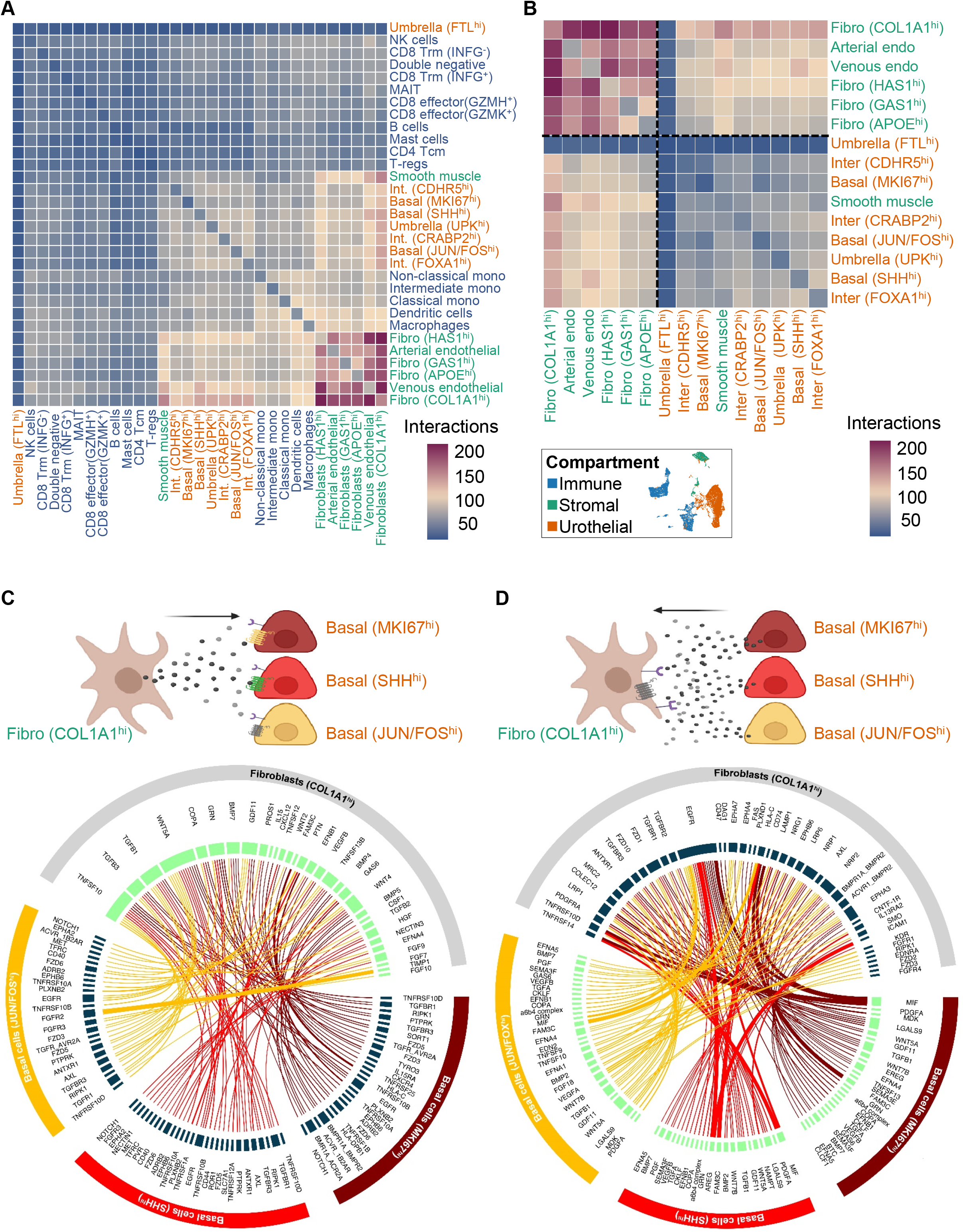
Identification of cell-cell communication networks in the adult human ureter. (A) & (B) Heatmaps of the number of predicted interactions between cell clusters across immune, stromal, and urothelial compartments (A) and between cells clusters in the stromal and urothelial compartments (B). (C) & (D) Circos plots showing non-integrin mediated interactions between *COL1A1^hi^* fibroblasts and three basal cell clusters, where ligands (green) are expressed in *COL1A1^hi^* fibroblasts and receptors (blue) are expressed in the basal cells in (C) and vice versa (D) The lines are color coded by basal clusters, and the thickness of lines represent the mean expression of the ligand and receptor in a given pair. Cell cartoons created with BioRender.com. See also Table S7.

Overall, the putative receptor-ligand communications in both directions showed significant enrichment for WNT, BMP, EGF, and TGFβ-related pathways (likelihood test, p value <0.00001). Such observation is consistent with a postulated importance of SHH expression in basal cells as SHH signaling induces stromal expression of WNT4 as well as the differentiation-associated BMP4 and BMP5 (Shin et al., 2011). Ligand expression in the *COL1A1*^hi^ fibroblast indicated this SHH-induced pattern (**Figure 5C**), with BMP4/5 signaling to the BMP receptors on the proliferative *MIK67*^hi^ basal cells, and WNT4 signaling to the frizzled class (FZD) receptors on all three basal cell populations. When we evaluated the reverse signaling from the basal cells to the fibroblasts, *WNT7B* ligand expression was detected in all three basal cell clusters and could signal to *FZD1* and *FZD10* expressed on *COL1A1*^hi^ fibroblasts (**Figure 5D**). In fact, studies in mice have pointed to an important role for the interaction between urothelial WNT7B and mesenchymal FZD1, working in parallel to SHH signaling, in ureter development (Jackson et al., 2020). The analysis also revealed Smoothened (*SMO*), another component of the SHH signaling network, to be expressed by the *COL1A1*^hi^ fibroblasts, with potential interactions with *BMP2* ligand from the *JUN/FOS*^hi^ and *SHH*^hi^ basal cell clusters. In addition, there were numerous predicted interactions from all three basal cell clusters with *EGFR* on the *COL1A1*^hi^ fibroblasts. Altogether, these results indicated that there are most likely numerous pathways working in parallel with *SHH* signaling network to mediate proliferation and differentiation of the adult ureter, and that the *COL1A1*^hi^ fibroblasts may be a central hub for mediating these processes.

### Sonic hedgehog expression predicts ureter organoid generation efficiency

Ureter organoids have emerged as a potential tool to study ureter development and homeostasis (Mullenders et al., 2019). To start laying the foundation for ureter cell culturing and differentiation *ex vivo*, we wanted to determine the fidelity of our cell cryopreservation method in maintaining cellular composition and gene expression patterns as well as to establish the ability of these cryopreserved cells to initiate organoid growth. To this end, we performed scRNAseq on freshly dissociated single cells and cryopreserved single cells from the same patient (U2), as well as the organoid derived from these same cryopreserved cells (**Figure 6A**). We detected the major cell types in the immune, stromal, and urothelial compartments as seen in the larger cohort analysis containing all ten human ureters (**Figures 1A**, **S5A**, and **S5B**). Based on scRNAseq results, we confirmed that the cryopreserved cells (Thawed) mirrored the freshly dissociated cells (Fresh) in terms of cellular composition (**Figures 6B** and **S5C**). The organoid, however, showed striking enrichment for basal and intermediate urothelial cell clusters and de-enrichment of the stromal and immune cell types (**Figures 6B** and **S5C**). The urothelial nature of the organoid and the enrichment for basal and intermediate cells were also confirmed by immunofluorescence staining of multiple ureter organoids (**Figure 6C**).

**Figure 6.**
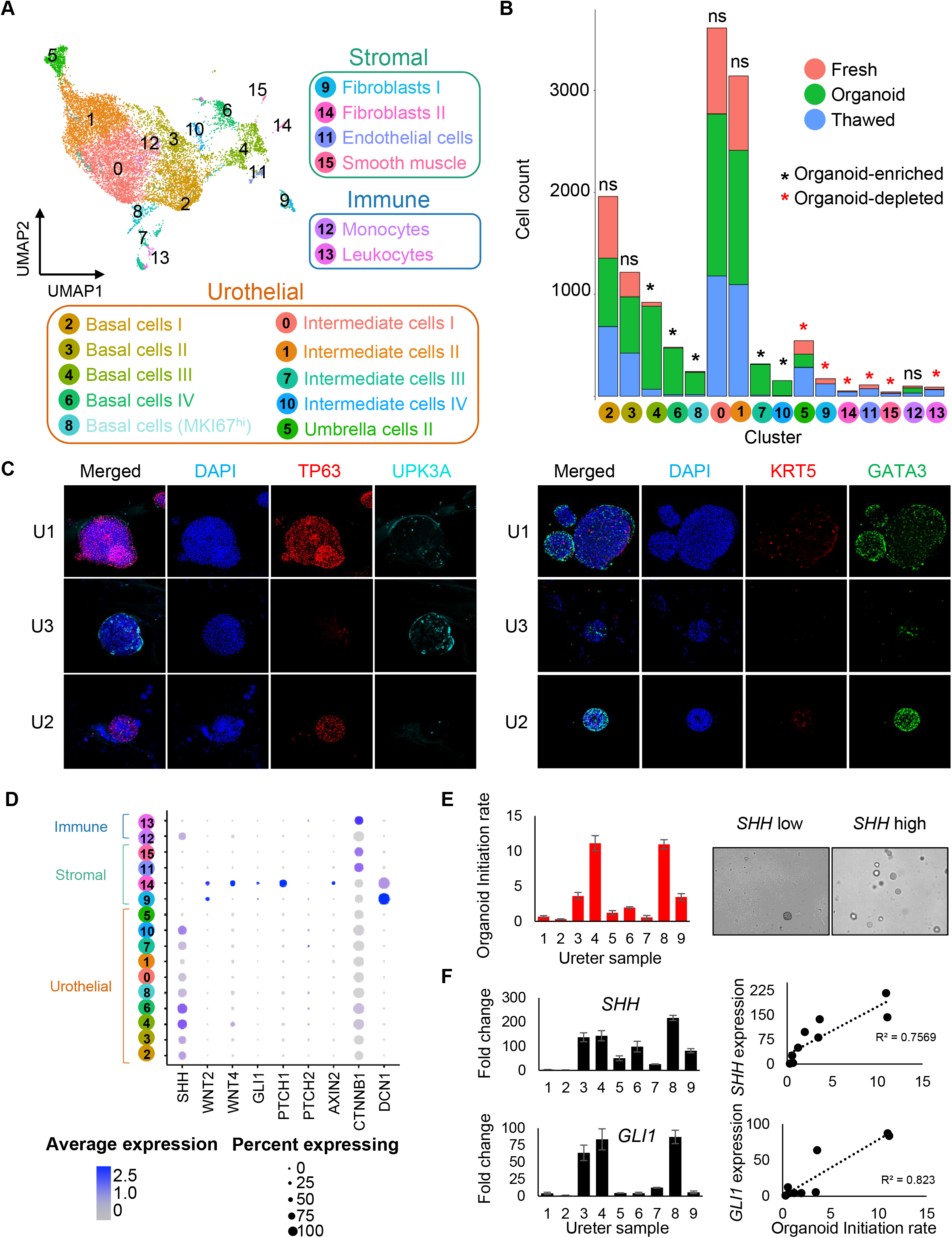
Sonic hedgehog expression predicts ureter organoid generation rate. (A) UMAP based on gene expression. Cells are colored by cluster. (B) Cell count within clusters stratified by sample: freshly processed (‘Fresh’), cryopreserved (‘Thawed’), and organoid. Clusters where the organoid sample showed statistically significant enrichment or depletion were marked by asterisks (FDR < 0.05 and absolute log2 fold change >1). (C) Immunofluorescence staining of ureter organoids. (D) Dotplot shows expression of Sonic Hedgehog pathway genes and Wnt pathway genes across cell clusters (E) Organoid initiation rate (right side) and light microscope images (left side). (F) RT-PCR expression (left) and correlation between RNA expression and organoid initiation rate (right). See also Figure S5.

Specific to the organoid, there was a notable expansion of the *MKI67*^hi^ basal cells (**Figure 6B**), as would be expected by the organoid culturing conditions that promote proliferation of adult stem cells (Mullenders et al., 2019). Yet, the exact identity of the urothelial self-renewing progenitor cells is still an area of intense debate (Colopy et al., 2014; Gandhi et al., 2013; Liu et al., 2019; Shin et al., 2011). Our results thus far implicated the *SHH*-expressing basal cells to have stem cell-like properties in adult human ureter (**Figure 4D**) and suggested that functional SHH signaling is necessary for proper tissue differentiation (**Figures 5C** and **5D**). Consistent with this, the organoid-enriched clusters (4, 6, 8, 7, & 10) all expressed *SHH*, with the highest levels observed in basal cell clusters 4 and 6 (**Figure 6D**). On the other hand, downstream *SHH* pathway genes, including *WNT4*, *GLI1*, and *PTCH1*, were robustly detected in cluster 14 fibroblasts, which contained the *COL1A1*^hi^ fibroblasts described in detail above (**Figure S5A**). Together, these observations led us to hypothesize that the organoid initiation rate would be positively correlated with the activity of the SHH pathway in individual patient specimens. To test this, we prospectively measured organoid initiation rates (**Figure 6E**) and quantified expression of *SHH* and its downstream targets for 9 additional patient samples (**Figures 6F** and **S5D**). Indeed, we observed that samples with higher levels of SHH signaling correlated with higher numbers of organoids formed (R^2^>0.7, **Figures 6F** and **S5D**). Moreover, this effect seemed to be specific as performing the same analysis with catenin beta 1 (*CTNNB1*) did not show such a correlation (R^2^=0.3535, **Figure S5D**). While the underlying reasons for intrinsic differences in *SHH* expression by sample was not investigated, our data supported the *SHH*-expressing basal cells having stem cell-like properties that can promote human ureter organoid formation *in vitro*.

## Discussion

Ureteral defects and injuries remain a significant clinical burden and, without successful repair, can greatly diminish the quality of life in patients affected by them. Tissue engineering offers a unique and promising approach to repairing these damages. Conventional tissue engineering approaches require the use of biomaterials that, ideally, can (1) preserve the anatomy and functionality of the tissue, (2) provide the appropriate structural and mechanical properties, and (3) recapitulate a microenvironment that can support proper differentiation of organ-specific cell types and secretion of their own extracellular matrix proteins that will eventually replace the artificial scaffold (Simaioforidis et al., 2013). The use of artificial scaffolds has some inherent drawbacks (Roy et al., 2020). For example, cells may fail to fully integrate into the matrix scaffold during bladder tissue reconstruction (Bouhout et al., 2016). Furthermore, interactions between the epithelia and stroma are required for the proper maturation of the urothelium (Shin et al., 2011; Wang et al., 2017). To avoid these pitfalls, stromal self-assembly methods, where tissue-derived stromal cells are utilized for producing cell type-specific extracellular matrix, have been proposed (Roy et al., 2020). The “self-assembly” technique has been used to generate various tissues (Saba et al., 2018), including a complete 3D human bladder (Bouhout et al., 2016). Although this approach is generally more time-consuming and costly, the benefits seem to outweigh the disadvantages. In order to implement a successful strategy for ureter tissue engineering, we must understand the populations of cells, their signaling networks, and their spatial relations within a healthy human ureter. Such a detailed understanding will not only guide the selection of methods used to generate the tissue but also help establish quality-control parameters to ensure that proper cellular composition and tissue architecture are achieved in the engineered tissue. These fundamental data about the human ureter are missing from our current knowledge base so we set out to generate a cellular atlas of the healthy human ureter.

Leveraging single-cell RNAseq, we analyzed 30,942 ureter cells, spanning the immune (16 clusters), stromal (7 clusters), and urothelial (8 clusters) compartments, to delineate the cellular constituents of the ureter. The rich diversity of immune cells in normal human ureters was revealed for the first time, highlighting a need to compare immune cell distributions from pathological conditions to understand how this organ defends itself. We also conducted the first ever spatial gene expression analysis of the human ureter and additional immunofluorescence assays to validate findings from scRNAseq and to spatially visualize previously unappreciated cell types within this organ, including four distinct types of fibroblasts. Interestingly, one of the fibroblast subtypes was characterized by high expression of *HAS1*. Hyaluronan-based biomaterials are a popular and emerging scaffold material in tissue engineering (Zamani et al., 2020). Thus, this fibroblast subtype could be central for the future development of tissue-engineered ureters. It also points towards the consideration of hyaluronan-based materials as a suitable scaffold. Moreover, our in-depth analysis of the stromal compartment may also help identify media formulations to stimulate the isolation and propagation of specific populations of patient-derived fibroblasts. This could expedite extracellular matrix deposition, which is a common obstacle for stromal self-assembly methods. For example, the *HAS1*^hi^ fibroblasts expressed high levels of *SEMA3C* (semaphorin 3C). Addition of recombinant SEMA3A, another member of the semaphorin family proteins, to organoid cultures has been shown to significantly increase organoid size (Karpus et al., 2019). Thus, including SEMA3C as an additive into the ureter organoid culture media could be of interest in developing self-assembled ureter tissues for grafting.

The urothelium in the upper urinary tract, including that of the ureter, is derived from the mesoderm while the bladder urothelium arises from the endoderm. However, most studies of the urothelium discount this intrinsic difference and consider the urothelium of the bladder and the ureter as a singular entity (Dalghi et al., 2020). As such, finer intricacies of cell types and expression patterns in the ureter have been historically overlooked (Dalghi et al., 2020; Liang et al., 2005; Wu et al., 2009). With this dataset, we had the opportunity to enumerate ureter-specific urothelial cell types and their respective gene expression patterns. For instance, we discovered a new population of umbrella cells, expressing high levels of *FLT*, *FTH1*, and *KRT20*. The existence of this population was independently confirmed by immunofluorescent staining of FLT and KRT20 protein. The role of ferritin in the ureter is not well studied; however, regulation of iron availability via sequestration by ferritin is a key host cell defense against uropathogens (Bauckman et al., 2019; Bauckman and Mysorekar, 2016). Excessive iron bioavailability in the bladder urothelium has been shown to increase uro-pathogenic growth (Bauckman and Mysorekar, 2016). Therefore, this cluster of *FTL^hi^* umbrella cells may have some unique roles in urothelial defense against pathogens by chelating excess iron in a non-bioavailable reservoir. It will be interesting to see if similar cells are also present in mice ureters and to test whether ablation might have functional consequences. Similarly, single cell analysis from patients with ureter infections may further implicate this umbrella cell subtype in urothelial defense. Finally, this rare population of terminally differentiated umbrella cells provides a unique benchmark for assessing success of engineered ureters.

Our study also showed that ureter urothelial lineage specification shares some common themes with that of the bladder, despite having different embryonic origin. Retinoic acid (RA) signaling (Gandhi et al., 2013), peroxisome proliferator-activated receptors (PPAR) signaling (Liu et al., 2019), and sonic hedgehog (SHH) signaling (Shin et al., 2011) have all been implicated in the bladder urothelium maintenance, specification, and differentiation. Retinoids are potent signaling molecules involved in urothelial differentiation in embryonic stem cells as well as the adult steady-state urothelium (Gandhi et al., 2013). Our data revealed one of the intermediate clusters to be defined by high levels of cellular retinoic acid binding protein 2 (*CRABP2*). CRABP2 facilitates transport of retinoic acids to retinoic acid receptors (RARs) to activate their activity. In addition, one of the basal-to-intermediate cell trajectories was enriched for genes, including retinol binding protein 2 (*RBP2*), in the “vitamin digestion and absorption” KEGG pathway. PPAR signaling, which has been shown to be an upstream regulator of mouse bladder urothelial differentiation during tissue repair, also emerged from our cell trajectory analysis as a potential driver of basal-to-intermediate cell differentiation in the human ureter. Our data also corroborated previous findings that SHH is a critical mediator of progenitor cell maintenance in the adult urothelium (Shin et al., 2011). We detected *SHH*-expressing basal cells in the human ureter and determined that these cells are critical for organoid formation *in vitro*.

We also identified epidermal growth factor receptor (EGFR) signaling as a potential new player in ureter urothelial specification. First, we detected two subtypes of basal cells that contained *SHH*-expressing cells, but one was concurrently characterized by high levels of *JUN* and *FOS* expressions. JUN and FOS together form the activator protein 1 (AP-1) complex. AP-1 is a downstream signaling module of the EGFR pathways and activates transcription of *AREG*, which further stimulates EGF receptor as part of a positive feedback loop (Frohlich et al., 2015). Second, EGFR receptor-ligand interactions were among the top predicted communications between *COL1A1*^hi^ fibroblasts and all basal cell subtypes. For example, EGFR expressed on the *COL1A1*^hi^ fibroblasts can potentially respond to basal cell-secreted MIF, COPA, TGFB, TGFA, AREG, GRN, and BTC. Taken together, these observations implicate a complex interaction between EGFR and SHH signaling axes in ureter homeostasis that warrants further research. Moreover, it will be important to delineate how each of the four signaling pathways (RA, PPAR, SHH, and EGFR) contributes to ureter repair in response to injuries and infections.

Beyond providing a cellular atlas of the human ureter and illuminating key directions for future mechanistic investigations, our study also generated knowledge that are directly relevant to ureter tissue engineering efforts. Based on our analyses, fibroblast subtypes in the human ureter have distinct expression profiles suggestive of their unique functions in supporting urothelial cell differentiation and growth. In particular, the *COL1A1^hi^* fibroblasts and the *HAS1^hi^* fibroblasts should be seriously considered in tissue engineering experiments. The *COL1A1^hi^* fibroblasts reside closest to the basal urothelial cells *in vivo* and may provide critical signaling inputs to support urothelial stem cell maintenance, growth, and differentiation. The *HAS1^hi^* fibroblasts, on the other hand, may be essential for creating a ureter-specific extracellular matrix (ECM) that ensures proper organ architecture. Inclusion of these patient-derived ureter fibroblasts in self-assembly methods could be pivotal to their success. Furthermore, our data can also be used to optimize tissue culture media that not only support stem cell growth but also promote appropriate differentiation to form a human ureter that is suitable for transplantation. Last but not least, the cell type markers identified in this study will help us isolate specific cell populations from patient tissues to initiate culturing, determine the correct proportions of target cell types to ensure growth, and develop quality-control assays for engineered ureters.

In conclusion, our studies offer a comprehensive analysis of the many cell populations within the human ureter, as well as the first spatial transcriptomic analysis of this tissue type. This work will provide an important reference for future research into directing the accurate engineering of ureters in regenerative medicine. In addition, this work will aid in our fundamental understanding of the normal physiology, transcriptional landscape, and signaling networks in the adult ureter, a tissue that has largely been overlooked.

## Acknowledgements

We thank the Lerner Research Institute Flow Cytometry Core and Genomics Core for 10x single cell experiment support, Computing Services for data and computing resource management, and Imaging Core for Visium spatial gene expression experiment support. We also acknowledge Jean R. Clemenceau for his early contribution to our single cell data analysis workflow. This work is supported by K08 CA237842 to B.H.L. and Cleveland Clinic institutional support to A.H.T.

## Author Contributions

Conceptualization, B.H.L. and A.H.T.; Methodology, E.E.F., S.S., B.H.L., and A.H.T.; Software, S.S. and M.B., Investigation, E.E.F., S.S., U.T., P.D., M.B., H.Q., M.W., and M.N.; Resources, M.E., A.W., G.H., and B.H.L.; Writing – Original Draft, E.E.F., S.S., and A.H.T.; Writing – Review & Editing, O.W. and B.H.L.; Visualization, E.E.F., S.S., O.W., and A.H.T.; Funding Acquisition, B.H.L. and G.H.; Supervision, O.W., B.H.L., and A.H.T.

## Declaration of interest

The authors declare no competing interests

## STAR Methods

### Lead Contact and Materials Availability

Correspondence and requests for materials should be addressed to the Lead Contact, Angela H. Ting. This study did not generate new unique reagents.

### Data and Code Availability

Raw and processed sequencing data have been deposited in the Gene Expression Omnibus under accession numbers GSE184111 (scRNA-seq) and GSE184112 (Visium spatial gene expression) and will be made public once the manuscript has been accepted for publication in a peer-reviewed journal. Microscopy data reported in this paper will be shared by the lead contact upon request. The analysis code and accompanying output files are available at (https://github.com/2019surbhi/tinglab_ureter_analysis/).

### Experimental Model and Subject Details

#### Human Ureter Procurement and Single Cell Isolation

Fresh human ureter tissues were collected at the Cleveland Clinic. Approval was received by the Institutional Review Board at Cleveland Clinic, and subsequently signed informed consent from all patients was received. Ten different samples were excised from patients undergoing radical cystectomy. The age and sex of the patients are described in Table S1. After cutting off blood supply, until the sample was removed from the body, the total amount of time was < 30 minutes. The fresh tissue sample was transferred from the operating room to the laboratory on ice, with the entire transport time being less than ten minutes. The tissues were immediately washed with sterile PBS and centrifuged for 5 minutes at 1000xg. The PBS was aspirated off and the tissues were transferred to a sterile petri dish, where they were minced using sterile razor blades until a “paste-like” consistency was achieved (< 5 minutes). The tissue digestion was carried out using the Papain Dissociation System (Worthington Biochemical Corporation) according to the manufactures instructions with, minor changes. Briefly, 5 mL of sterile Earle’s Balanced Salt Solution (EBSS) was added to one vial of papain and warmed at 37°C in a water bath. 500 μL EBSS was added to one vial of DNase and placed on ice. The minced tissue was transferred to a 50 mL conical containing 5 mL papain and 500 μL DNase. The tissue was incubated with shaking (70rpm) at 37°C for 1hr. 5.5 mL of ovoid inhibitor was added to the tissue suspension to stop the digestion. Large debris was removed by passing the entire volume (~11 mL) through a 70 μm cap filter. The cell suspension was subsequently centrifuged at 1000 rpm for 5 min to pellet cells, and the supernatant was discarded. The cell pellet was re-suspended in 3 mL ACK lysis buffer (Thermo Fisher Scientific), incubated at RT for 3 min, centrifuged at 1000 rpm for 5 min, and washed with 5 mL PBS. The cell pellet was re-suspended in an appropriate volume of PBS based on its size. The re-suspended cells were then passed through a 40 μm Flowmi tip (Bel-Art). Cells were counted with trypan blue to note cell viability. The target concentration was 1,000 cells/μL.

### Method Details

#### Sample Processing with 10X Genomics and cDNA Library Preparation

We used 10X Genomics Chromium Single Cell 3’ Reagents Kit version 3.1. We targeted to recover 10,000 cells per sample unless a given sample had fewer than 10,000 total cells. All samples had >5,000 cells captured for library generation and sequencing. Following the 10X protocol, we added the single cell suspension, the gel beads, and the emulsion oil to the 10X Genomics Single Cell Chip G and ran the Chromium Controller. Immediately following the droplet generation, samples were transferred to a PCR 8-tube strip (USA scientific), and reverse transcription was performed using SimpliAmp thermal cycler (Applied Biosystems). Following reverse transcription, cDNA was recovered using the recovery reagent provided by 10X Genomics. The cDNA was cleaned up using the Silane DynaBeads according to the 10X Genomics user guide. The purified cDNA was amplified for 11 cycles and subsequently cleaned up using SPRIselect beads (Beckman Coulter). To determine the cDNA concentrations, 1:10 dilution of each sample was analyzed on an Agilent Bioanalyzer High Sensitivity chip. The cDNA libraries were constructed according to the Chromium Single Cell 3’ Reagent Kit version 3.1 user guide.

#### scRNA data analysis

The scRNA-seq libraries were pooled and sequenced to a target depth of 50,000 read pairs per cell. De-multiplexed FASTQ files were processed using cellranger-4.0.0, where reads were mapped using the count pipeline with the pre-built reference genome refdata-gex-GRCh38-2020A and GTF from GENECODE v32 (GRCh28.p13). The downstream analysis was performed using Seurat 3.2.1 (Stuart et al., 2019) in R. The raw data, comprising of 64,350 cells, was filtered for low-quality cells using QC thresholds determined by assessing the distribution plots. Briefly, cells with mitochondrial reads >25% of total mapped reads, gene counts < 200 or >8,000, and total mapped read counts <2,000 were filtered out, leaving 30,942 high-quality cells for the downstream analysis. Seurat’s standard workflow was followed. Using the 2,000 most variable genes, principal components (PCs) were computed, and the first 50 PCs were utilized to generate clustering at resolution 0.5. Cells were into UMAPs for visualization, and cell clusters were annotated by marker gene expression. Cluster level similarity was assessed by calculating cluster correlations (Pearson’s) based on the top 2000 variable genes, which grouped the clusters into 3 major compartments – urothelial, stromal, and immune. Subset analysis for each compartment was performed using the same Seurat workflow. Clustering optimization was performed using clustree (Zappia and Oshlack, 2018) and silhouette plots (Kassambara and Mundt, 2020; Yan and Marr, 2005).

The sequencing data from fresh, thawed, and organoid samples derived from a single patient (U2) were analyzed using the same pipeline. Cells with mitochondrial reads >30% of total mapped reads, gene counts < 250 or >8,000, and total mapped read counts <7,000 were filtered out, yielding 13,161 high-quality cells for further analysis. Clustering was performed using the first 30 PCs at resolution 0.3. Cell types enriched in organoid sample were determined using Permutation Test as described in https://github.com/rpolicastro/scProportionTest at a threshold of absolute log2 fold difference >1.

#### scRNA subset analysis

For each subset analysis, cells from clusters assigned to a given compartment (immune, stromal, or urothelial) were processed by the same scRNA analysis pipeline as described above, starting from raw gene counts. The subsets were subjected to additional filtering and re-processing until any cross-compartment contamination was sufficiently removed. Potentially contaminating clusters were removed ‘*If the top100 differential marker list of a given cluster shows markers that should be exclusively expressed by cells of another compartment* (e.g. PTPRC appears among top100 markers in a stromal subset cluster)’ **AND** ‘*the contaminating marker is expressed in >10% of cells in that cluster*’. Additionally, clustree tool was utilized to visualize median gene expression pattern of the contaminating marker(s) to determine if a certain clustering parameter (number of PC and clustering resolution) effectively clusters out these contaminating cells. If such parameters exist, the subset was clustered using these parameters and subjected to the same subset analysis workflow, while excluding the contaminating cluster(s). Finally, each subset data was clustered using the first 50 PCs at resolution 0.5, with the exception of the urothelial subset (resolution 0.2).

#### Monocle analysis

The urothelial subset cells were subjected to trajectory analysis using the standard workflow of Monocle3 (Cao et al., 2019; Qiu et al., 2017; Trapnell et al., 2014). Briefly, the raw gene count per cell was log normalized and scaled prior to computing principal components (PCs). First 50 PCs were used for batch correction. The UMAP coordinates from Seurat analysis were transferred to the monocle object to preserve the cluster identities across the two analysis sets. Monocle3 built a trajectory across the cells, based on gene expression changes that signify the different cell states transitions within the dataset. It created convergent and divergent branches in the trajectory (when setting loop =TRUE) to denote multiple possible outcomes of cell state transitions. Next, cells across the trajectory were ordered based on user defined starting point (root) for the trajectory. The lowest Pseudotime score was assigned to the root (cells with the highest *SHH* expression, which are in cluster 3) while the remaining cells were assigned a score based on the tool’s perception of subsequent order of the cell state transitions. Genes that vary as a function of Pseudotime were deduced by running graph_test function and plotted on Pseudotime scale.

#### Fibroblast gene set enrichment analysis

Differential gene list was generated for the 4 Fibroblast clusters in the stromal subset using Seurat’s FindMarkers function. The upregulated markers were filtered in each list using a threshold of log fold change (log FC) >0.58, which is equivalent to log2FC 1.5. Gene Ontology (GO) enrichment analysis for each cluster gene list and cluster-wide comparison was performed using clusterProfiler (Yu et al., 2012).

#### CellPhoneDB analysis

Normalized gene counts for stromal and urothelial scRNA data were utilized to determine significant ligand-receptor interactions using CellPhoneDB (Efremova et al., 2020) with p value cutoff as 0.0001. Non-integrin ligand-receptor pairs involving secreted factors were assessed for signaling between *COL1A1^hi^* fibroblasts (cluster 5 in the stromal subset) and the three basal cell subtypes (clusters 2, 3, and 6 in the urothelial subset). A circos plot described in iTALK (Wang et al., 2019) was adapted to plot the interactions between the *COL1A1^hi^* fibroblasts and the individual basal cell clusters.

#### Visium spatial gene expression library generation

Ureter samples were collected immediately upon surgical removal, embedded in Optimal Cutting Temperature (OCT) compound, and stored in −80°C until ready to process. Per the Visium Spatial Tissue Optimization User Guide (CG000238 Rev D), 6 minutes was determined to be the optimum permeabilization time for the ureter tissue. Sections of 10 μm thickness were placed on Visium Spatial Gene Expression Slide. Tissue fixation and H&E staining were performed following the 10X Genomics Tissue Preparation Guide. Bright field images were taken with a Slide Scanning Microscope-Helios (Leica Microsystems) at 10x magnification. Libraries were prepared according to the Visium Spatial Gene Expression User Guide with the following parameters: 6min permeabilization time, 16 cycles for cDNA amplification, and 16 cycles for the sample index PCR. Libraries were pooled and sequenced to a target depth of 50,000 read pairs per tissue covered spot on Capture Area.

#### Visium data analysis

De-multiplexed FASTQ files were processed by the 10X spaceranger software (v1.3.0). The “count” pipeline was executed using the pre-built GRCh38-2020A reference genome. For transcriptomics annotation, we used GTF from GENECODE v32 (GRCh38.p13). Spaceranger was configured with the “reorient-images” option, so that the optimal fiducial alignment could be achieved by the software. Differential gene expression analysis was then processed using in R (v4.0.3) using Seurat (v4.0.3) (Hao et al., 2021). First, the percentage of mitochondrial reads was identified for each Visium spot. The data was then normalized using SCTransform (Hafemeister and Satija, 2019). This method identifies variance on a spot-by-spot basis using a negative binomial regression model, and is intended to conserve greater biological significance than log normalization methods. SCTransform was run with the “vars.to.regress” option set to regress out the percentage of mitochondrial reads in each spot, as performed in the vignette for SCTransform (Hafemeister and Satija, 2019). Figure generation was done via the Seurat::SpatialFeaturePlot function, run entirely with default options for each feature included. Plots were saved with ggsave individually in png format with 300 dpi per image.

#### Ureter organoid generation and SHH gene expression

Following tissue dissociation of ureters, one aliquot of cells were counted, and 1,000 cells were re-suspended in 15 μl matrigel (Corning) per bead at 4°C for seeding into one well of a 24-well plate. Once matrigel solidified, 0.5 mL of urothelial growth media (Mullenders et al., 2019) was added to each well. Media was refreshed every other day. The number of organoids initiated after two weeks was counted and initiation rate was calculated as the number of organoids formed per original number of single cells seeded per well. A second aliquot of each dissociated sample was set aside following tissue dissociation for RNA isolation using the AllPrep DNA/RNA Kit (Qiagen). Total RNA was converted to cDNA using SuperScript™ VILO™ cDNA Synthesis Kit (Thermo Fisher). Following cDNA conversion, each qRT-PCR reaction was carried out with QuantiTect SYBR green reagent (Qiagen) on the Roche LightCycler® 96. PCR primers for specific SHH pathway genes can be found the **Key Resources Table**. Results were based on biological triplicates that were assayed by technical duplicate PCR reactions, and statistical significance was determined by Student’s t test

#### Immunofluorescence imaging

Human ureters or organoids were fixed in 4% paraformaldehyde overnight at 4°C. Ureters were embedded in paraplast, sectioned at 10 μm, dewaxed, and processed for citrate-based antigen retrieval before the incubation with the primary antibodies. Species-specific Alexa-488, −555, and −647 secondary antibodies (Invitrogen) were used for visualization. In the cases where antibodies of the same species had to be used for co-staining, Tyramide Signal Amplification (Biotium) was used for the first primary antibody followed by antibody stripping and incubation with the second primary antibody. Slides were embedded in Prolong Diamond (Invitrogen) and imaged on a Leica DM5500 using the LASX software and image deconvolution. The following primary antibodies were used: APOE (Sigma); COL15A1 (Sigma); Cytokeratin 5 (Cell Signaling); Cytokeratin 17 (Invitrogen); Cytokeratin 20 (Invitrogen); Ferritin Light Chain (Invitrogen); FOXF1 (R&D); GATA3 (Cell Signaling); MKI67 (Cell Signaling); LIF (Sigma); LYPD3 (Invitrogen); P63 (Cell Signaling); Smooth Muscle Actin (Sigma); Uroplakin 3A (LS Biosciences).

## Supplemental material

**Figure S1.**
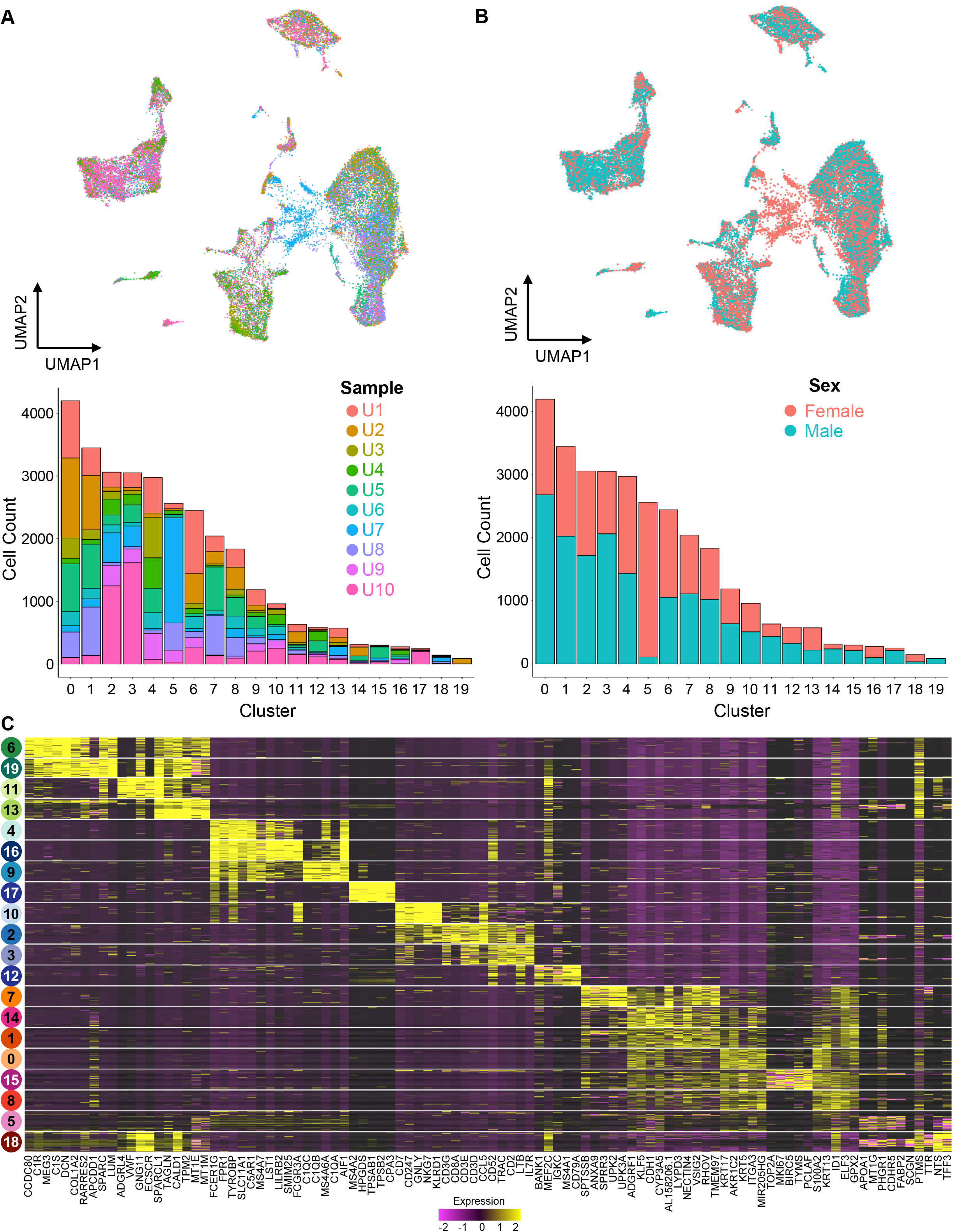
UMAP based on gene expression (top) and barplots corresponding to number of cells per cluster (bottom). Cells are colored by (A) sample and (B) Sex. (C) Heatmap for top 5 differential markers for each cluster.

**Figure S2.**
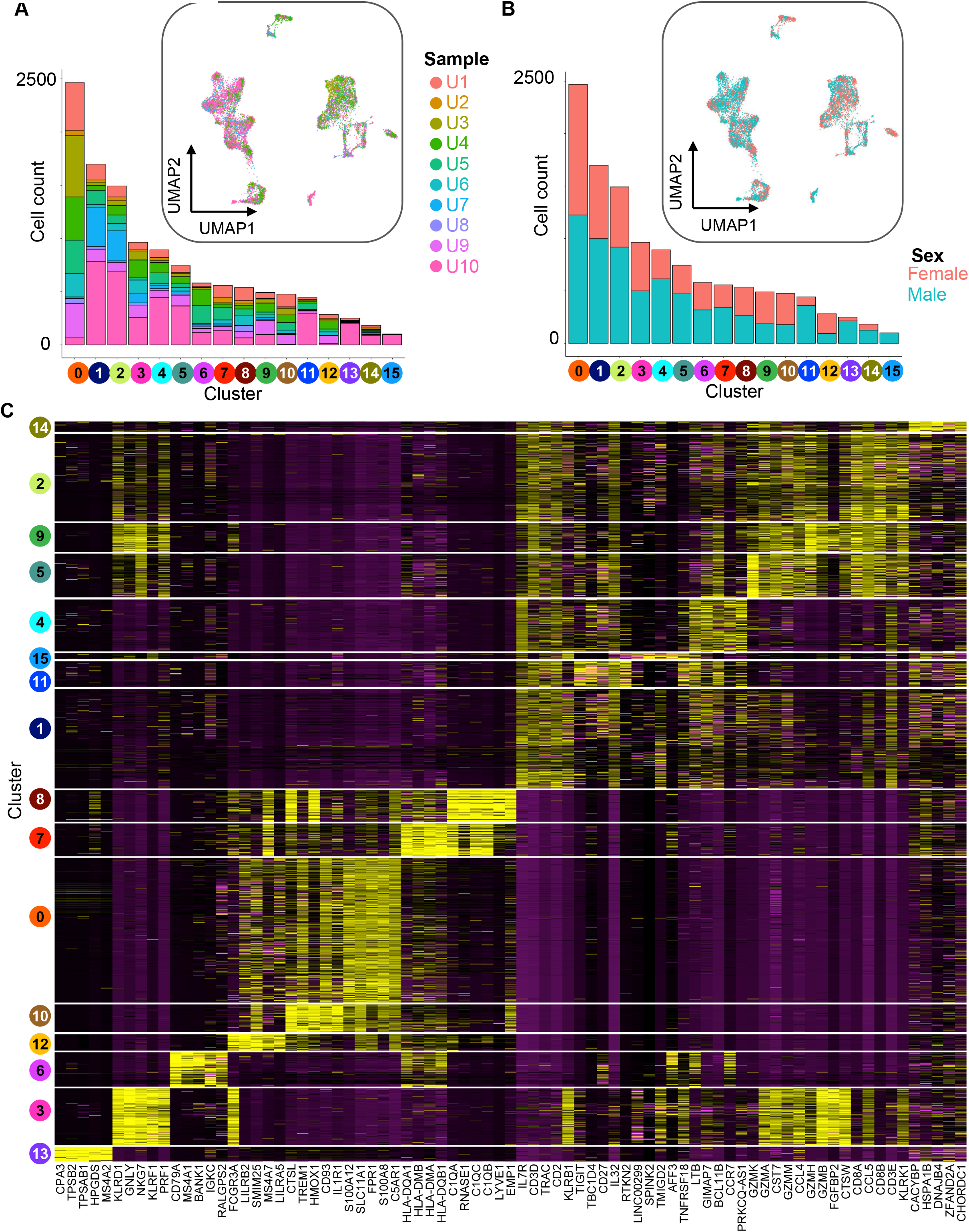
UMAP based on gene expression (top) and barplots corresponding to number of cells per cluster (bottom). Cells are colored by (A) sample and (B) Sex. (C) Heatmap for top 5 differential markers for each cluster.

**Figure S3.**
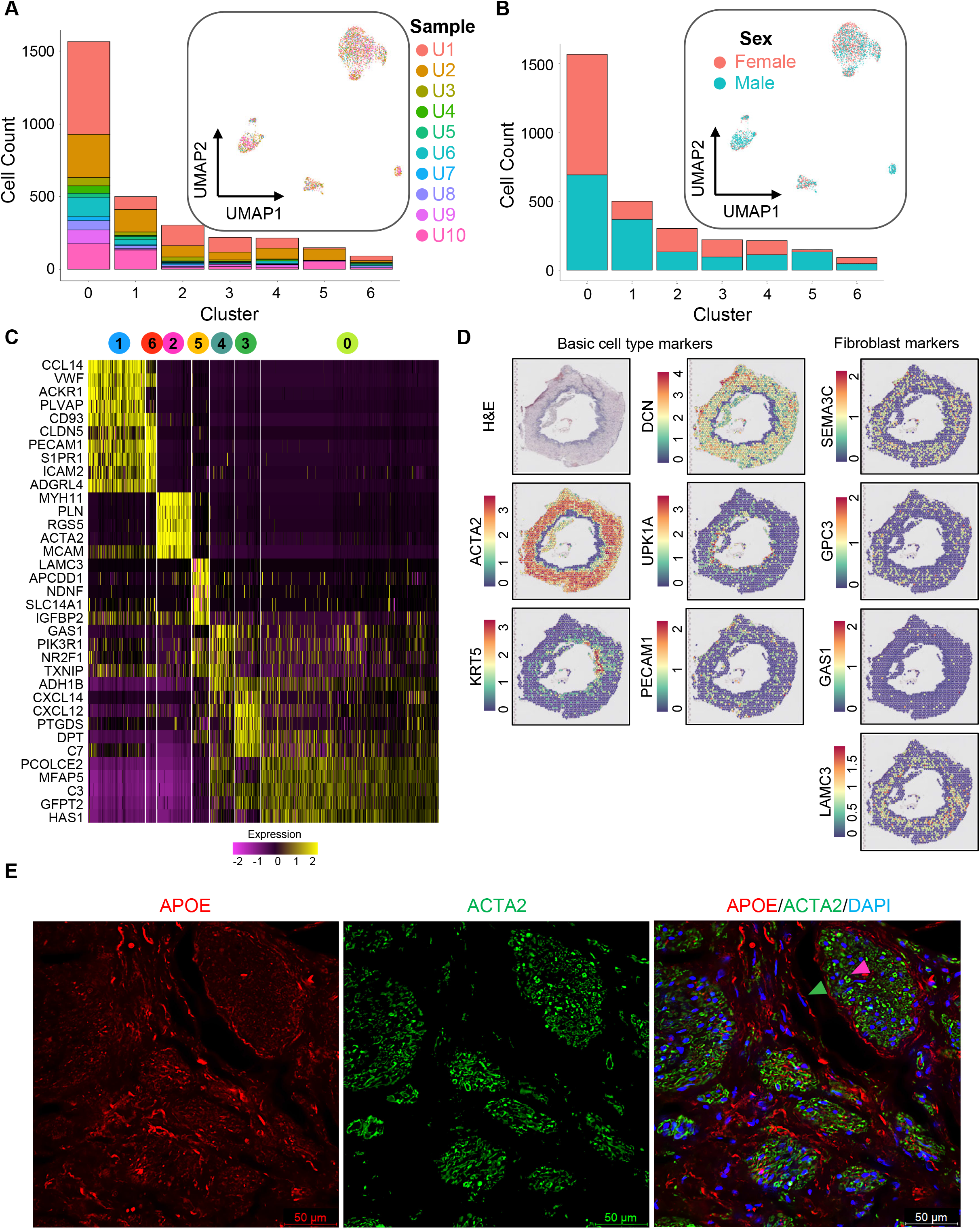
UMAP based on gene expression (top) and barplots corresponding to number of cells per cluster (bottom). Cells are colored by (A) sample and (B) Sex. (C) Heatmap for top 5 differential markers for each cluster. (D) Visium spatial gene expression in log-normalized (logNorm) counts for selected genes in human ureter cross-sections. (E) Immunofluorescence staining of a human ureter cross-section. Arrows are color-coded by cluster.

**Figure S4.**
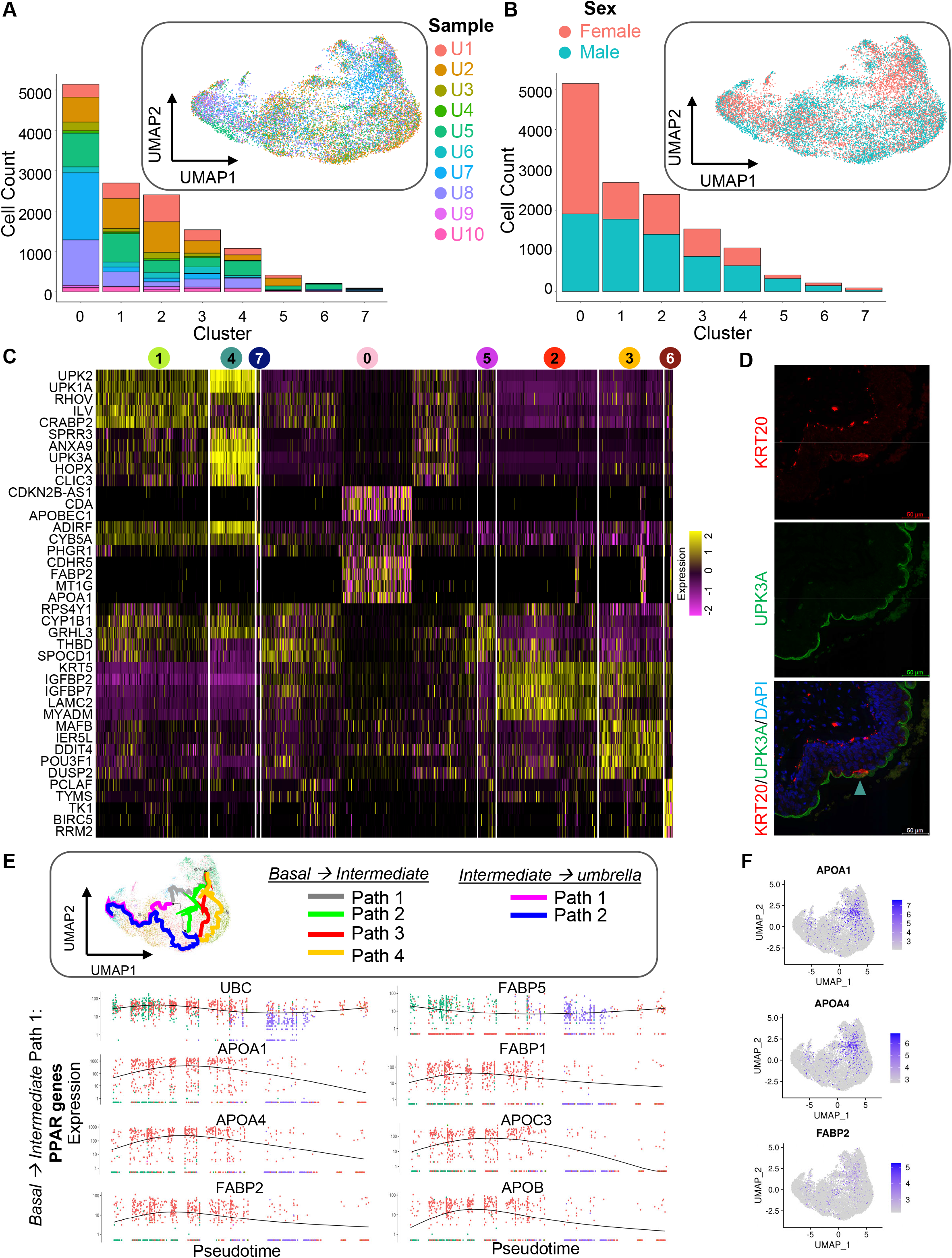
UMAP projections and barplots for urothelial subset with cells colored by (A) sample and (B) Sex. (C) Heatmap for top 5 differential markers for each cluster. (D) Immunofluorescence staining of a human ureter cross-section. Arrows are color-coded by cluster. (E) Monocle analysis on urothelial subset (top). Expression pattern across the pseudotime trajectory for select genes from PPAR signaling (bottom). (F) Feature plots for urothelial subset of select PPAR signaling genes with scale bar indicating log normalized gene counts.

**Figure S5.**
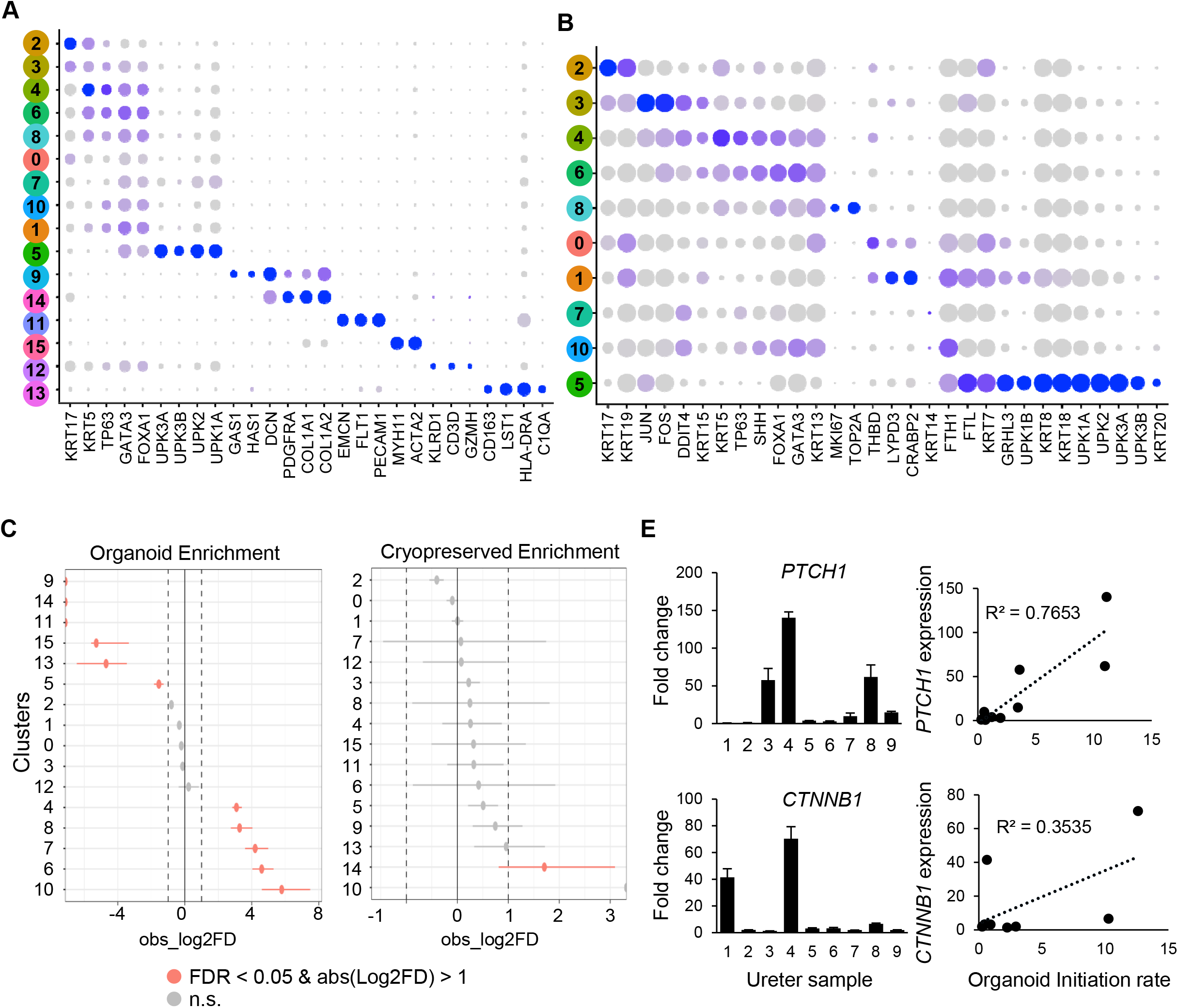
(A) & (B) Dot plots show the expression level of known markers in (A) all clusters and (B) urothelial clusters. (C) Statistical comparison of cell proportions between organoid cells and non-organoid cells (Organoid Enrichment, left) or between cryopreserved (thawed) cells and freshly isolated (fresh) cells (Cryopreserved Enrichment, right) within each cluster using scProportion function. Both tests were performed for log2FD > 1 (log2 fold difference between a given comparison - organoid vs non organoid or fresh vs thawed) and FDR < 0.05. Cell proportions that are not significantly (n.s.) different for a given cluster-wise comparison are colored in grey. The lines denote the confidence interval. The out-of-bound values (±∞) for log2FD are shown as partial circles. (D) RT-PCR expression (left) and correlation between RNA expression and organoid initiation rate (right).

**Table S1. Summary statists of samples and sequencing data (Related to Figure 1).**

**Table S2. Global ureter cluster differentially expressed genes (Related to Figure 1).**

**Table S3. Immune subset cluster differentially expressed genes (Related to Figure 2).**

**Table S4. Stromal subset cluster differentially expressed genes (Related to Figure 3.**

**Table S5. Urothelial subset cluster differentially expressed genes (Related to Figure 4).**

**Table S6. KEGG enrichment on dynamic genes from cell trajectory analysis (Related to Figure 4).**

**Table S7. CellphoneDB analysis output (Related to Figure 5).**

## Notes

### Competing Interest Statement

The authors have declared no competing interest.

## References

Armand, E.J., Li, J., Xie, F., Luo, C., and Mukamel, E.A. (2021). Single-Cell Sequencing of Brain Cell Transcriptomes and Epigenomes. Neuron 109, 11–26.

Bauckman, K.A., Matsuda, R., Higgins, C.B., DeBosch, B.J., Wang, C., and Mysorekar, I.U. (2019). Dietary restriction of iron availability attenuates UPEC pathogenesis in a mouse model of urinary tract infection. Am J Physiol Renal Physiol 316, F814–F822.

Bauckman, K.A., and Mysorekar, I.U. (2016). Ferritinophagy drives uropathogenic Escherichia coli persistence in bladder epithelial cells. Autophagy 12, 850–863.

Bohnenpoll, T., Feraric, S., Nattkemper, M., Weiss, A.C., Rudat, C., Meuser, M., Trowe, M.O., and Kispert, A. (2017a). Diversification of Cell Lineages in Ureter Development. J Am Soc Nephrol 28, 1792–1801.

Bohnenpoll, T., Wittern, A.B., Mamo, T.M., Weiss, A.C., Rudat, C., Kleppa, M.J., Schuster-Gossler, K., Wojahn, I., Ludtke, T.H., Trowe, M.O., et al. (2017b). A SHH-FOXF1-BMP4 signaling axis regulating growth and differentiation of epithelial and mesenchymal tissues in ureter development. PLoS Genet 13, e1006951.

Bouhout, S., Chabaud, S., and Bolduc, S. (2016). Organ-specific matrix self-assembled by mesenchymal cells improves the normal urothelial differentiation in vitro. World J Urol 34, 121–130.

Cao, J., Spielmann, M., Qiu, X., Huang, X., Ibrahim, D.M., Hill, A.J., Zhang, F., Mundlos, S., Christiansen, L., Steemers, F.J., et al. (2019). The single-cell transcriptional landscape of mammalian organogenesis. Nature 566, 496–502.

Chen, Z., Zhou, L., Liu, L., Hou, Y., Xiong, M., Yang, Y., Hu, J., and Chen, K. (2020). Single-cell RNA sequencing highlights the role of inflammatory cancer-associated fibroblasts in bladder urothelial carcinoma. Nat Commun 11, 5077.

Colopy, S.A., Bjorling, D.E., Mulligan, W.A., and Bushman, W. (2014). A population of progenitor cells in the basal and intermediate layers of the murine bladder urothelium contributes to urothelial development and regeneration. Dev Dyn 243, 988–998.

Combes, A.N., Phipson, B., Lawlor, K.T., Dorison, A., Patrick, R., Zappia, L., Harvey, R.P., Oshlack, A., and Little, M.H. (2019). Single cell analysis of the developing mouse kidney provides deeper insight into marker gene expression and ligand-receptor crosstalk. Development 146.

Cui, H.J., Cui, X.G., Jing, X., Yuan, Y., Chen, Y.Q., Sun, Y.X., Zhao, W., and Liu, X.G. (2019). GAS1 Deficient Enhances UPR Activity in Saccharomyces cerevisiae. Biomed Res Int 2019, 1238581.

Dalghi, M.G., Montalbetti, N., Carattino, M.D., and Apodaca, G. (2020). The Urothelium: Life in a Liquid Environment. Physiol Rev 100, 1621–1705.

DeGraff, D.J., Clark, P.E., Cates, J.M., Yamashita, H., Robinson, V.L., Yu, X., Smolkin, M.E., Chang, S.S., Cookson, M.S., Herrick, M.K., et al. (2012). Loss of the urothelial differentiation marker FOXA1 is associated with high grade, late stage bladder cancer and increased tumor proliferation. PLoS One 7, e36669.

dela Paz, N.G., and D’Amore, P.A. (2009). Arterial versus venous endothelial cells. Cell Tissue Res 335, 5–16.

Efremova, M., Vento-Tormo, M., Teichmann, S.A., and Vento-Tormo, R. (2020). CellPhoneDB: inferring cell-cell communication from combined expression of multi-subunit ligand-receptor complexes. Nat Protoc 15, 1484–1506.

England, A.R., Chaney, C.P., Das, A., Patel, M., Malewska, A., Armendariz, D., Hon, G.C., Strand, D.W., Drake, K.A., and Carroll, T.J. (2020). Identification and characterization of cellular heterogeneity within the developing renal interstitium. Development 147.

Evanko, S.P., Potter-Perigo, S., Petty, L.J., Workman, G.A., and Wight, T.N. (2015). Hyaluronan Controls the Deposition of Fibronectin and Collagen and Modulates TGF-beta1 Induction of Lung Myofibroblasts. Matrix Biol 42, 74–92.

Frohlich, H., Bahamondez, G., Gotschel, F., and Korf, U. (2015). Dynamic Bayesian Network Modeling of the Interplay between EGFR and Hedgehog Signaling. PLoS One 10, e0142646.

Gandhi, D., Molotkov, A., Batourina, E., Schneider, K., Dan, H., Reiley, M., Laufer, E., Metzger, D., Liang, F., Liao, Y., et al. (2013). Retinoid signaling in progenitors controls specification and regeneration of the urothelium. Dev Cell 26, 469–482.

Gild, P., Kluth, L.A., Vetterlein, M.W., Engel, O., Chun, F.K.H., and Fisch, M. (2018). Adult iatrogenic ureteral injury and stricture-incidence and treatment strategies. Asian J Urol 5, 101–106.

Godfrey, D.I., Koay, H.F., McCluskey, J., and Gherardin, N.A. (2019). The biology and functional importance of MAIT cells. Nat Immunol 20, 1110–1128.

Hafemeister, C., and Satija, R. (2019). Normalization and variance stabilization of single-cell RNA-seq data using regularized negative binomial regression. Genome Biol 20, 296.

Hao, Y., Hao, S., Andersen-Nissen, E., Mauck, W.M., 3rd, Zheng, S., Butler, A., Lee, M.J., Wilk, A.J., Darby, C., Zager, M., et al. (2021). Integrated analysis of multimodal single-cell data. Cell 184, 3573–3587 e3529.

Hayes, B.W., and Abraham, S.N. (2016). Innate Immune Responses to Bladder Infection. Microbiol Spectr 4.

Hu, Y., Hu, Y., Xiao, Y., Wen, F., Zhang, S., Liang, D., Su, L., Deng, Y., Luo, J., Ou, J., et al. (2020). Genetic landscape and autoimmunity of monocytes in developing Vogt-Koyanagi-Harada disease. Proc Natl Acad Sci U S A 117, 25712–25721.

Jackson, A.R., Ching, C.B., McHugh, K.M., and Becknell, B. (2020). Roles for urothelium in normal and aberrant urinary tract development. Nat Rev Urol 17, 459–468.

Jiang, D., Liang, J., and Noble, P.W. (2007). Hyaluronan in tissue injury and repair. Annu Rev Cell Dev Biol 23, 435–461.

Karpus, O.N., Westendorp, B.F., Vermeulen, J.L.M., Meisner, S., Koster, J., Muncan, V., Wildenberg, M.E., and van den Brink, G.R. (2019). Colonic CD90+ Crypt Fibroblasts Secrete Semaphorins to Support Epithelial Growth. Cell Rep 26, 3698–3708 e3695.

Kassambara, A., and Mundt, F. (2020). factoextra: Extract and Visualize the Results of Multivariate Data Analyses. R package version 1.0.7.

Lake, B.B., Chen, S., Hoshi, M., Plongthongkum, N., Salamon, D., Knoten, A., Vijayan, A., Venkatesh, R., Kim, E.H., Gao, D., et al. (2019). A single-nucleus RNA-sequencing pipeline to decipher the molecular anatomy and pathophysiology of human kidneys. Nat Commun 10, 2832.

Li, Y., Xu, X., Song, L., Hou, Y., Li, Z., Tsang, S., Li, F., Im, K.M., Wu, K., Wu, H., et al. (2012). Single-cell sequencing analysis characterizes common and cell-lineage-specific mutations in a muscle-invasive bladder cancer. Gigascience 1, 12.

Liang, F.X., Bosland, M.C., Huang, H., Romih, R., Baptiste, S., Deng, F.M., Wu, X.R., Shapiro, E., and Sun, T.T. (2005). Cellular basis of urothelial squamous metaplasia: roles of lineage heterogeneity and cell replacement. J Cell Biol 171, 835–844.

Liu, C., Tate, T., Batourina, E., Truschel, S.T., Potter, S., Adam, M., Xiang, T., Picard, M., Reiley, M., Schneider, K., et al. (2019). Pparg promotes differentiation and regulates mitochondrial gene expression in bladder epithelial cells. Nat Commun 10, 4589.

Loperena, R., Van Beusecum, J.P., Itani, H.A., Engel, N., Laroumanie, F., Xiao, L., Elijovich, F., Laffer, C.L., Gnecco, J.S., Noonan, J., et al. (2018). Hypertension and increased endothelial mechanical stretch promote monocyte differentiation and activation: roles of STAT3, interleukin 6 and hydrogen peroxide. Cardiovasc Res 114, 1547–1563.

Menon, R., Otto, E.A., Kokoruda, A., Zhou, J., Zhang, Z., Yoon, E., Chen, Y.C., Troyanskaya, O., Spence, J.R., Kretzler, M., et al. (2018). Single-cell analysis of progenitor cell dynamics and lineage specification in the human fetal kidney. Development 145.

Mullenders, J., de Jongh, E., Brousali, A., Roosen, M., Blom, J.P.A., Begthel, H., Korving, J., Jonges, T., Kranenburg, O., Meijer, R., et al. (2019). Mouse and human urothelial cancer organoids: A tool for bladder cancer research. Proc Natl Acad Sci U S A 116, 4567–4574.

Paffenholz, P., and Heidenreich, A. (2021). Modern surgical strategies in the management of complex ureteral strictures. Curr Opin Urol 31, 170–176.

Qiu, X., Mao, Q., Tang, Y., Wang, L., Chawla, R., Pliner, H.A., and Trapnell, C. (2017). Reversed graph embedding resolves complex single-cell trajectories. Nat Methods 14, 979–982.

Rehman, S., and Ahmed, D. (2021). Embryology, Kidney, Bladder, and Ureter. In StatPearls (Treasure Island (FL)).

Riedel, I., Liang, F.X., Deng, F.M., Tu, L., Kreibich, G., Wu, X.R., Sun, T.T., Hergt, M., and Moll, R. (2005). Urothelial umbrella cells of human ureter are heterogeneous with respect to their uroplakin composition: different degrees of urothelial maturity in ureter and bladder? Eur J Cell Biol 84, 393–405.

Roy, V., Magne, B., Vaillancourt-Audet, M., Blais, M., Chabaud, S., Grammond, E., Piquet, L., Fradette, J., Laverdiere, I., Moulin, V.J., et al. (2020). Human Organ-Specific 3D Cancer Models Produced by the Stromal Self-Assembly Method of Tissue Engineering for the Study of Solid Tumors. Biomed Res Int 2020, 6051210.

Saba, I., Jakubowska, W., Bolduc, S., and Chabaud, S. (2018). Engineering Tissues without the Use of a Synthetic Scaffold: A Twenty-Year History of the Self-Assembly Method. Biomed Res Int 2018, 5684679.

Sheng, X., Guan, Y., Ma, Z., Wu, J., Liu, H., Qiu, C., Vitale, S., Miao, Z., Seasock, M.J., Palmer, M., et al. (2021). Mapping the genetic architecture of human traits to cell types in the kidney identifies mechanisms of disease and potential treatments. Nat Genet 53, 1322–1333.

Shin, K., Lee, J., Guo, N., Kim, J., Lim, A., Qu, L., Mysorekar, I.U., and Beachy, P.A. (2011). Hedgehog/Wnt feedback supports regenerative proliferation of epithelial stem cells in bladder. Nature 472, 110–114.

Simaioforidis, V., de Jonge, P., Sloff, M., Oosterwijk, E., Geutjes, P., and Feitz, W.F. (2013). Ureteral tissue engineering: where are we and how to proceed? Tissue Eng Part B Rev 19, 413–419.

Sole-Boldo, L., Raddatz, G., Schutz, S., Mallm, J.P., Rippe, K., Lonsdorf, A.S., Rodriguez-Paredes, M., and Lyko, F. (2020). Single-cell transcriptomes of the human skin reveal age-related loss of fibroblast priming. Commun Biol 3, 188.

Stuart, T., Butler, A., Hoffman, P., Hafemeister, C., Papalexi, E., Mauck, W.M., 3rd, Hao, Y., Stoeckius, M., Smibert, P., and Satija, R. (2019). Comprehensive Integration of Single-Cell Data. Cell 177, 1888–1902 e1821.

Teh, Y.C., Ding, J.L., Ng, L.G., and Chong, S.Z. (2019). Capturing the Fantastic Voyage of Monocytes Through Time and Space. Front Immunol 10, 834.

Terpstra, M.L., Remmerswaal, E.B.M., van Aalderen, M.C., Wever, J.J., Sinnige, M.J., van der Bom-Baylon, N.D., Bemelman, F.J., and Geerlings, S.E. (2020). Circulating mucosal-associated invariant T cells in subjects with recurrent urinary tract infections are functionally impaired. Immun Inflamm Dis 8, 80–92.

Trapnell, C., Cacchiarelli, D., Grimsby, J., Pokharel, P., Li, S., Morse, M., Lennon, N.J., Livak, K.J., Mikkelsen, T.S., and Rinn, J.L. (2014). The dynamics and regulators of cell fate decisions are revealed by pseudotemporal ordering of single cells. Nat Biotechnol 32, 381–386.

Volkmer, J.P., Sahoo, D., Chin, R.K., Ho, P.L., Tang, C., Kurtova, A.V., Willingham, S.B., Pazhanisamy, S.K., Contreras-Trujillo, H., Storm, T.A., et al. (2012). Three differentiation states risk-stratify bladder cancer into distinct subtypes. Proc Natl Acad Sci U S A 109, 2078–2083.

Vorobev, V., Beloborodov, V., Golub, I., Frolov, A., Kelchevskaya, E., Tsoktoev, D., and Maksikova, T. (2021). Urinary System Iatrogenic Injuries: Problem Review. Urol Int 105, 460–469.

Wang, C., Ross, W.T., and Mysorekar, I.U. (2017). Urothelial generation and regeneration in development, injury, and cancer. Dev Dyn 246, 336–343.

Wang, Y., Wang, R., Zhang, S., Song, S., Jiang, C., Han, G., Wang, M., Ajani, J., Futreal, A., and Wang, L. (2019). iTALK: an R Package to Characterize and Illustrate Intercellular Communication. bioRxiv, 507871.

Wolfien, M., David, R., and Galow, A.M. (2021). Single-Cell RNA Sequencing Procedures and Data Analysis. In Bioinformatics, I.N. Helder, ed. (Brisbane (AU)).

Wu, H., Malone, A.F., Donnelly, E.L., Kirita, Y., Uchimura, K., Ramakrishnan, S.M., Gaut, J.P., and Humphreys, B.D. (2018). Single-Cell Transcriptomics of a Human Kidney Allograft Biopsy Specimen Defines a Diverse Inflammatory Response. J Am Soc Nephrol 29, 2069–2080.

Wu, X.R., Kong, X.P., Pellicer, A., Kreibich, G., and Sun, T.T. (2009). Uroplakins in urothelial biology, function, and disease. Kidney Int 75, 1153–1165.

Yan, J., and Marr, T.G. (2005). Computational analysis of 3’-ends of ESTs shows four classes of alternative polyadenylation in human, mouse, and rat. Genome Res 15, 369–375.

Yang, Z., Li, C., Fan, Z., Liu, H., Zhang, X., Cai, Z., Xu, L., Luo, J., Huang, Y., He, L., et al. (2017). Single-cell Sequencing Reveals Variants in ARID1A, GPRC5A and MLL2 Driving Self-renewal of Human Bladder Cancer Stem Cells. Eur Urol 71, 8–12.

Yu, G., Wang, L.G., Han, Y., and He, Q.Y. (2012). clusterProfiler: an R package for comparing biological themes among gene clusters. OMICS 16, 284–287.

Yu, Z., Liao, J., Chen, Y., Zou, C., Zhang, H., Cheng, J., Liu, D., Li, T., Zhang, Q., Li, J., et al. (2019). Single-Cell Transcriptomic Map of the Human and Mouse Bladders. J Am Soc Nephrol 30, 2159–2176.

Yu, Z., Mannik, J., Soto, A., Lin, K.K., and Andersen, B. (2009). The epidermal differentiation-associated Grainyhead gene Get1/Grhl3 also regulates urothelial differentiation. EMBO J 28, 1890–1903.

Zamani, F., Zare Shahneh, F., Aghebati-Maleki, L., and Baradaran, B. (2013). Induction of CD14 Expression and Differentiation to Monocytes or Mature Macrophages in Promyelocytic Cell Lines: New Approach. Adv Pharm Bull 3, 329–332.

Zamani, M., Shakhssalim, N., Ramakrishna, S., and Naji, M. (2020). Electrospinning: Application and Prospects for Urologic Tissue Engineering. Front Bioeng Biotechnol 8, 579925.

Zappia, L., and Oshlack, A. (2018). Clustering trees: a visualization for evaluating clusterings at multiple resolutions. Gigascience 7.

